# Unraveling the *Gardnerella* Pangenome: Insights into Genetic Diversity, Virulence, and Antibiotic Resistance

**DOI:** 10.1101/2025.03.03.641253

**Authors:** Pedro Teixeira, Márcia Sousa, Teresa Gonçalves Ribeiro, Filipa Grosso, Luísa Peixe

## Abstract

In recent years, the development of user-friendly and accessible bioinformatic tools has significantly advanced *Gardnerella* genome analysis. Despite the taxonomic reclassification of the genus in 2019, many studies continue to rely on subgroups/ clades for species identification. This leads to frequent misidentifications, hindering accurate assessments of species-specific features, such as antimicrobial resistance and pathogenic potential. This study aims to address these gaps by employing precise species-level identification to improve our understanding on *Gardnerella* and its clinical implications. We performed the most extensive pangenome analysis of *Gardnerella* to date, analyzing 153 genomes (120 from NCBI and 33 newly sequenced).

*G. vaginalis, G. leopoldii, G. swidsinskii,* and *G. pickettii* species exhibited open pangenomes. The *Gardnerella* pangenome is highly diverse, with 90% of its genes belonging to the accessory genome. Carbohydrate transport and metabolism contributed the most for the primary genomic differences observed among *Gardnerella* species. Additionally, the search for antibiotic resistance genes (ARGs) revealed *lsaC* gene as the most prevalent ARG, potentially conferring resistance to clindamycin, a commonly prescribed antibiotic for treating bacterial vaginosis (BV). The clustering of *Gardnerella* genomes mimics the core genome phylogeny, pointing to species specific virulence-associated factors, highlighting important factors such as vaginolysin and sialidases.

Our comprehensive pangenome analysis shows that *Gardnerella* species are well-adapted within the urogenital tract, with identified virulence factors and metabolic capabilities potentially enhancing the pathogenicity of some strains. Future studies should explore *Gardnerella*’s ecological interactions and pathogenic mechanisms within the urogenital microbiome.

**Importance:** In recent years, the availability of user-friendly and accessible bioinformatics tools has led to a surge in studies analyzing the *Gardnerella* genome. Despite the redefinition of the genus in 2019, many researchers continue to rely on subgroups or clades for *Gardnerella* identification. Our study builds upon the updated taxonomy of *Gardnerella*, offering a more comprehensive understanding of its genetic diversity. By conducting a pangenome analysis of 153 *Gardnerella* genomes, we uncover significant genetic variability, species-specific virulence factors, and the widespread presence of antibiotic resistance genes, highlighting the urgent need for precise species-level identification. The identification of novel genomic features and antibiotic resistance profiles not only enhances our understanding of *Gardnerella*’s clinical implications but also sheds light on its metabolic and virulence potential. These insights help clarify the pathogenic mechanisms of *Gardnerella* and provide a more refined understanding of its role in the urogenital microbiome.

## Background

Historically, *Gardnerella vaginalis* was considered the unique species within the genus. However, taxonomic revision in 2019 led to the emended description of *G. vaginalis* and the identification of three additional species: *Gardnerella leopoldii*, *Gardnerella piotii*, and *Gardnerella swidsinskii*, along with nine distinct *Gardnerella* genomic species (GG) (1–3). Further studies expanded this diversity with the recognition of an additional GG (4). More recently, our research group formally described two GG, *Gardnerella pickettii* (formerly GG3) and *Gardnerella greenwoodii* (formerly GG8) (2).

*Gardnerella*, particularly *G. vaginalis*, has long been associated with bacterial vaginosis (BV), a common vaginal dysbiosis linked to adverse pregnancy outcomes and increased susceptibility to sexually transmitted infections (5–8). However, the discovery of new *Gardnerella* species challenges the long-held assumption that *G. vaginalis* is the only species involved in BV and other urogenital infections. Moreover, *Gardnerella* species, including *G. vaginalis*, have been isolated from the urogenital tract, rectum, and midstream urine of asymptomatic women (9–12), highlighting their potential role as commensals. It is believed that BV development is triggered by the formation of polymicrobial biofilms, facilitated by the action of sialidases and the secretion of vaginolysin (13, 14). Notably, not all *Gardnerella* strains, including some *G. vaginalis* strains are associated with BV, exhibit sialidase activity nor encode vaginolysin (1, 15–17) Recent studies have also identified novel sialidases and vaginolysin variants (1, 15–17), underscoring the need for further investigation into their roles in BV pathogenesis.

Pangenome analysis provides valuable insights into the bacterial genetic diversity, evolution, and functional traits (4, 18, 19). This approach has been applied in *Gardnerella* research to explore species delineation, dynamics of lateral gene flow, metabolic capabilities, virulence factors (VFs), and evolutionary history (4, 18–23). However, some earlier studies relied on outdated taxonomic classifications (e.g., groups or clades encompassing different *Gardnerella* species) or misidentified isolates (24–26), emphasizing the need for caution when interpreting previous findings.

The increasing availability of *Gardnerella* genomes presents an opportunity to refine pangenome analysis by integrating updated taxonomic frameworks. A comprehensive genomic approach can provide a deeper understanding of the genetic factors and mechanisms that shape the diversity and pathogenic potential of *Gardnerella* species.

In this study, we present the most extensive *Gardnerella* genomic analyses to date, incorporating the latest taxonomic updates. Our research explores the phylogenetic relationships, resistome, and virulome of *Gardnerella*, providing novel insights into species-specific traits and potential pathogenicity. Additionally, we investigate the colonisation patterns and emphasise the importance of host health status in understanding the complex interplay between *Gardnerella* and the urogenital tract.

## Results

### Overview of the *Gardnerella* Genomes Collection

A total of 153 *Gardnerella* genomes were analysed, including 120 publicly available genomes (15 previously unexplored) and additional 33 newly sequenced genomes from our collection (Table S1). Most genomes were available in draft form, reflecting the current state of *Gardnerella* genome assembly. The majority of the *Gardnerella* strains analysed (59.7%, n=89) originated from vaginal samples, with the remaining strains primarily sourced from urine samples (38.3%, n=57). We assessed the correlation between *Gardnerella* strains and human health conditions, identifying BV (38.3%, n=57) and overactive bladder syndrome (12.1%, n=18) as the most prevalent conditions, while 27.5% (n=41) of isolates originated from asymptomatic human. Unfortunately, a subset of *Gardnerella* isolates (18.1%, n=27) catalogued in the National Center for Biotechnology Information (NCBI) database lacked information regarding the human health status.

### Genomic Analyses Reveal Frequent Misidentification of *Gardnerella* sp

Approximately 57% of *Gardnerella* genomes in the NCBI Refseq database are misidentified or classified as *Gardnerella* sp. or *G. vaginalis*, likely due to their deposition before the formal description of new *Gardnerella* species (1, 2, 4). We employed a comprehensive genomic approach, combining Average Nucleotide Identity (ANI), digital DNA-DNA hybridization (dDDH), and core genome phylogeny to delineate the species and infer the phylogenetic relationships among the *Gardnerella* strains.

ANI and dDDH network analysis grouped *Gardnerella* strains into 16 clusters, adhering to established species delineation thresholds (70% for dDDH and 96% for ANI) (27–29) (Figure 1). Two previously unreported genomic clusters were identified: a potential new genomic species (GG15), comprising strains N165 and JCP7275, and GG16, represented by strain JNFY17 (Figure 1). However, core-genome phylogenetic analysis of 92 single-copy genes repositioned these strains within GG2 and *G. pickettii*, respectively. Moreover, the phylogenomic analysis supported a classification of 6 species and 8 GG, consistent with previous research (Figure 2) (4, 20).

**Figure 1.**
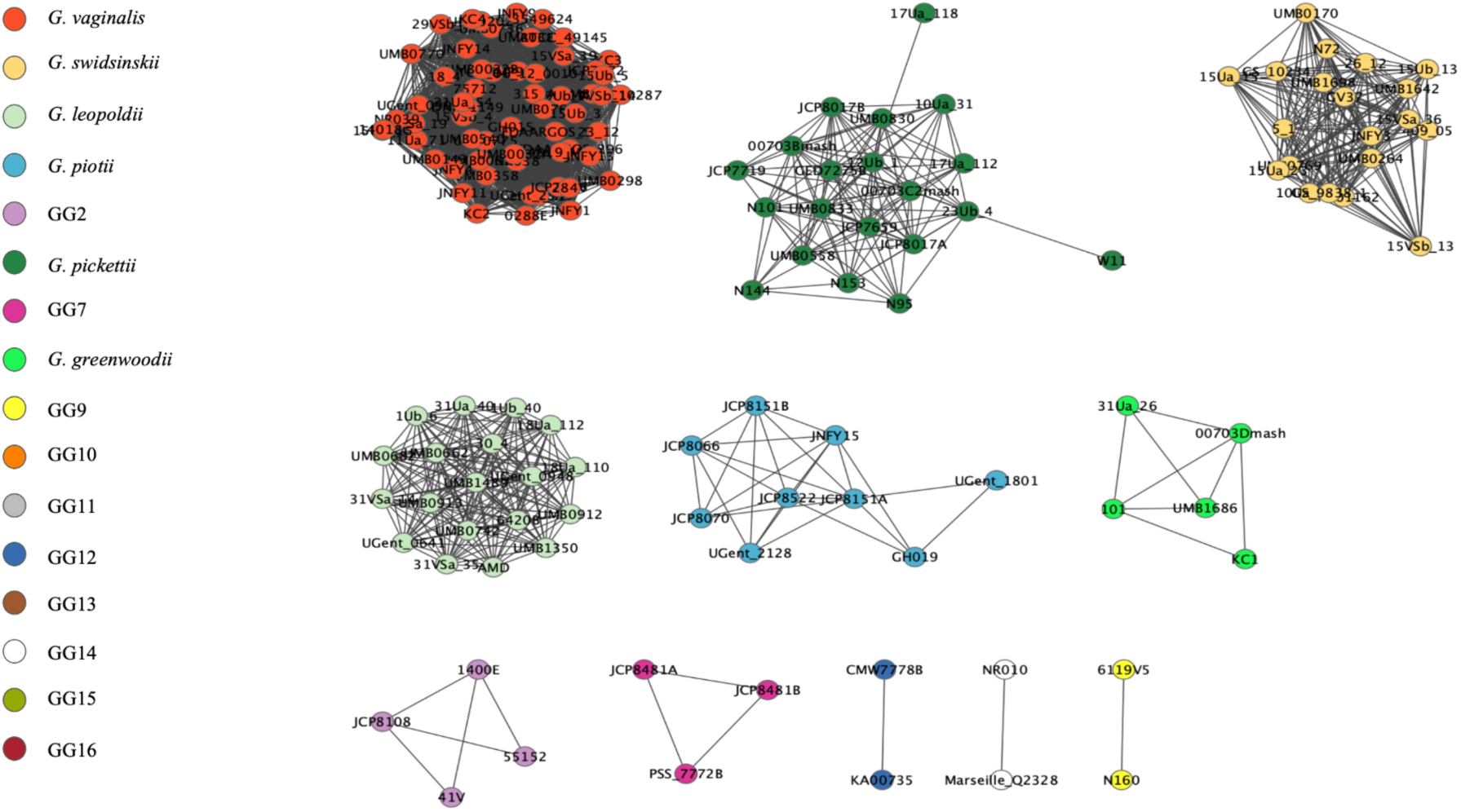
ANI- and dDDH-based network analysis of *Gardnerella* genomes. Each node represents a *Gardnerella* isolate, and node colors correspond to different species. Nodes connected by edges indicate clusters with ANI and dDDH values exceeding 96% and 70%, respectively.

**Figure 2.**
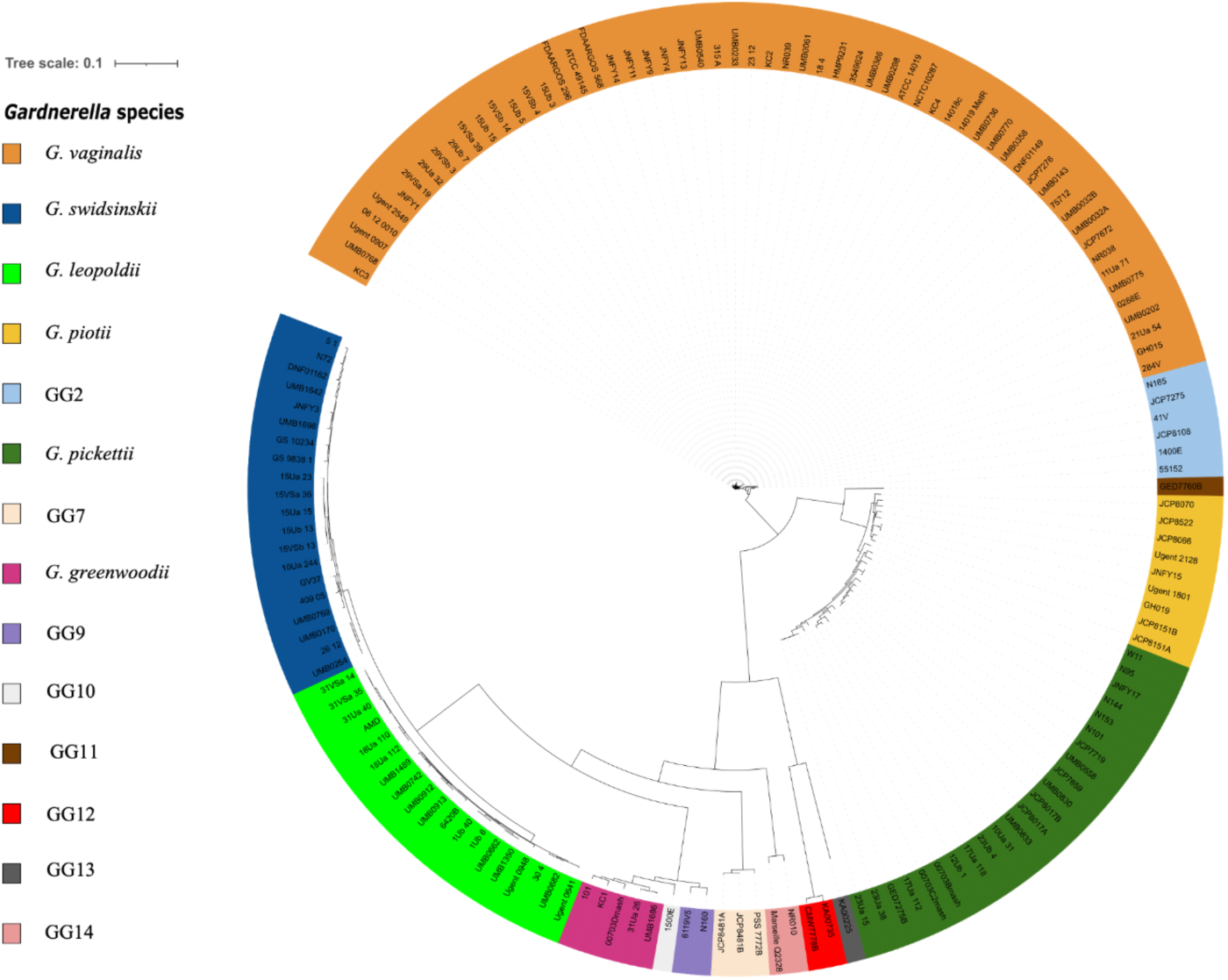
Core genome maximum likelihood phylogeny of *Gardnerella* genomes based on the alignment of the concatenation of amino acid sequences from the 92 single-copy core genes using anvi’o.

Among the identified species, *G. vaginalis* stands out as the most prevalent (38.5%, n=59), followed by *G. pickettii* (15.0%, n=23), *G. leopoldii* (13.1%, n=20), *G. swidsinskii* (12.4%, n=19), *G. piotii* (5.9%, n=9), and *G. greenwoodii* (3.3%, n=5). The remaining GG accounted for 11.8% (n=18) of the dataset (Figure 2, Table S1).

The genome size varied across *Gardnerella* species, ranging from 1.50 ± 0.02 Mbp (GG9) to 1.69 ± 0.06 Mbp (*G. vaginalis*) (p < 2.2x10^-16^), with an overall average of 1.61 ± 0.09 Mbp (Figure S1). Guanine-Cytosine (GC) content also differed, with GG12 exhibiting the lowest (38.1 ± 0.1%) and GG14 the highest (45.4 ± 0.1%) (p < 2.2x10^-16^) (Figure S2). Additionally, the number of coding sequences (CDS) ranged from 1,124 ± 0.7 (GG9) to 1,287 ± 61 (*G. vaginalis*) (p < 6.8x10⁻¹⁰) (Figure S3).

### Genomic Plasticity in *Gardnerella*: Evidence of Open Pangenomes

The pangenome of *Gardnerella* genus encompasses a total of 4824 gene clusters, categorised into 505 core genes (present in 100% of the genomes), 316 soft core genes (present in > 95% of genomes), 2584 dispensable genes (present in more than two genomes but less than 95%), and 1419 strain-specific genes (singletons). The core genome specifically includes 92 single-copy core genes shared across all 153 assemblies (Table S2). Approximately 10% of the pangenome is represented by the core genome, while the remaining 90% comprises accessory genome, including soft core, dispensable and singleton genes.

Variations in sample sizes across *Gardnerella* species can introduce bias when determining whether a pangenome is open or closed. To address this, we conducted species-specific pangenome analyses for *G. vaginalis*, *G. leopoldii*, *G. swidsinskii*, and *G. pickettii*. These analyses confirmed that all four species exhibit open pangenomes (Figure S4).

Based on core phylogeny and SNP matrix analysis, we identified identical clonal strains present in distinct biological samples from the same donor, specifically in urine and vaginal swabs, across *G. vaginalis*, *G. leopoldii, G. piotii,* and *G. swidsinskii* species (Table S3) (Figure S5A-D). Notably, a *G. pickettii* clone differing by only one SNP was identified in donors with different health statuses from separate studies. Strain GED7275B was isolated from the vaginal swab of an asymptomatic woman, while strain c17Ua_112^T^ was recovered from the urine of a woman diagnosed with overactive bladder.

Following the identification of the *Gardnerella* core and accessory genomes, gene clusters underwent functional annotation. Out of the total 4841 gene clusters, only 40% were associated with known Clusters of Orthologous Groups of proteins (COG functions), and 26% were linked with Kyoto Encyclopedia of Genes and Genomes (KEGG) Orthology (KO).

### Functional Insights into Core and Accessory Genomes of *Gardnerella* Species

Metabolic functions predominated within both the core and accessory genomes of *Gardnerella*. In the core genome, proteins were primarily linked to translation, ribosomal structure, and biogenesis (19.6%), followed by carbohydrate (9.8%) and amino acid (9.7%) transport and metabolism. In the accessory genome, the most prevalent COG category was carbohydrate transport and metabolism (G), accounting for 9.7%, with categories J (translation, ribosomal structure, and biogenesis) and L (replication, recombination, and repair) closely following at 7.3% each.

Further investigation into the functional categories of proteins encoded by isolates from different *Gardnerella* species revealed distinct clustering patterns. Non-metric multidimensional scaling (NMDS) analysis revealed a clear separation between *G. vaginalis* and GG2, as compared to other *Gardnerella* species (Figure 3).

**Figure 3.**
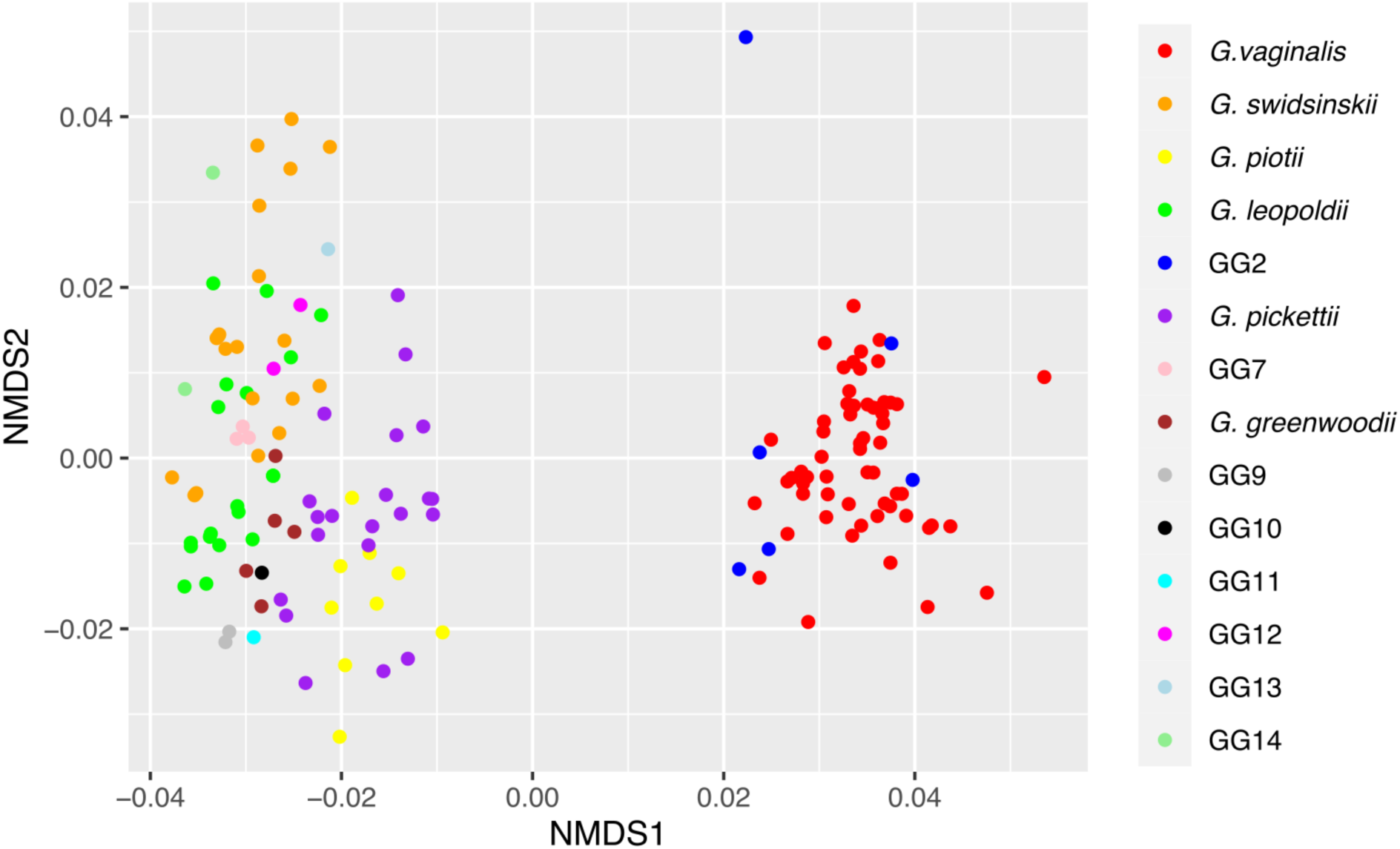
NMDS plot of *Gardnerella* spp. based on the distribution of COG categories.

The dissimilarity among species was explored through Similarity Percentage (SIMPER) analysis, which identifies the taxa that contribute most to the observed differences among sample groups. Along with *G. vaginalis* and GG2, *G. piotii* and *G. pickettii* also exhibited distinct grouping, separate from *G. leopoldii, G. swidsinskii*, and other GG (Figure 3). This differentiation was partly driven by the lower content of proteins linked to coenzyme transport and metabolism in *G. piotii* and *G. pickettii*, with a mean dissimilarity of 9.5 ± 1.4% (Figure 4A). The most prominent distinction between *G. vaginalis* and GG2, and the other *Gardnerella* species and GG was the proportion of proteins related to carbohydrate transport and metabolism, accounting for a mean dissimilarity of 32.1 ± 3.2% (Figure 4A). Additionally, the proportion of proteins associated with translation, ribosomal structure, and biogenesis was notably lower in *G. vaginalis* and GG2 compared to the other species (Figure 4B).

**Figure 4.**
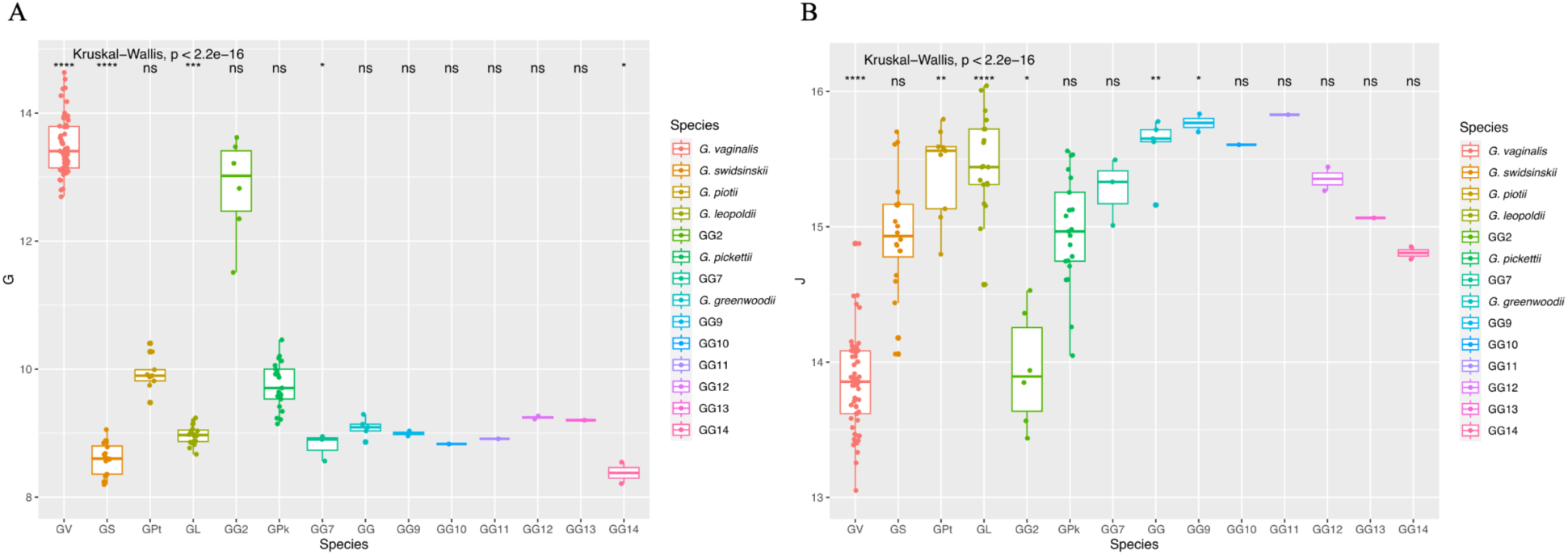
Differential abundance of key COG categories driving NMDS separation among *Gardnerella* spp. based on SIMPER analysis: **A)** Carbohydrate transport and metabolism (category G) and **B)** translation, ribosomal structure and biogenesis (category J).

Functional enrichment analysis revealed 13 KEGG modules and 3 COG pathways that were enriched in specific *Gardnerella* species (Table S4). Purine degradation, tryptophan, and asparagine biosynthesis were enriched in *G. leopoldii* and *G. swidsinskii*. Thiamine and molybdopterin biosynthesis were enriched in *G. leopoldii*, *G. swidsinskii*, and *G. vaginalis*, with only *G. leopoldii* and *G. swidsinskii* exhibiting molybdopterin adenylyltransferases (Table S4). The glucuronate and pentose phosphate pathways were enriched in *G. vaginalis*. Fatty acid biosynthesis, galactose degradation, and the non-phosphorylated Entner-Doudoroff pathway were enriched in *G. vaginalis*, *G. piotii*, and *G. pickettii*. These three species also shared enriched in functions related to lysine and menaquinone biosynthesis, keratan sulfate and methionine degradation, and glycolysis.

Notably, methionine degradation was enriched in *G. swidsinskii*, as well as *G. vaginalis*, *G. pickettii*, and *G. piotii*, due to the presence of 14 gene clusters encoding DNA-cytosine methylases. Of note, the functional enrichment analysis did not reveal significant differentiation based on isolation source or host disease associations of the different *Gardnerella* strains.

### Prevalence of *lsaC* Gene and Species Clustering based on Virulence Factor Profiles

We identified six antibiotic resistance genes (ARGs) in the pangenome: *aph(3’)-Ia* (encoding resistance to aminoglycosides), *tetM* (tetracyclines), *mefA* (macrolides), *msrD* (macrolides and streptogramin B), and *ermX* (macrolides, lincosamides, and streptogramins) and *lsaC* (macrolides, lincosamides, streptogramins A, and pleuromutilins) (Table S5). Among these, *lsaC*-like gene was most prevalent, present in 120 out of 153 genomes across all *Gardnerella* species, except for GG7 and GG13. The *mefA*-like and *msrD*-like genes were detected in all *Gardnerella* species (17 and 27 genomes, respectively), except GG9, GG13, and GG14. The *ermX* gene was found in only six strains (4 *G. vaginalis,* 1 *G. piotii*, and 1 *G. swidsinskii*), all isolated from BV-positive samples. The *tetM*-like gene was identified in 19 *Gardnerella* strains (8 *G. vaginalis*, 4 *G. pickettii*, 2 *G. swidsinskii*, 2 GG2, 1 *G. piotii*, 1 GG12, and 1 GG14), isolated from humans with different health conditions (pregnant woman, BV-positive women, diagnosed with overactive bladder).

We detected 117 different virulence-associated functions distributed across 261 gene clusters in the *Gardnerella* pangenome (Table S6). The clustering of *Gardnerella* genomes based on the presence/absence of virulence gene clusters mirrored the core genome phylogeny, pointing that virulence functions are species-specific within the genus (Figure S6). Two main clades emerged in the species clustering analysis: one composed of *G. vaginalis*, *G. piotii*, GG2, *G. pickettii* and GG11, and the other of the remaining *Gardnerella* species/GG. The differences between these two clusters were largely attributed to distinct virulence factor profiles, particularly those associated to adhesion, mucin degradation and immune evasion (Figure S6). A more detailed representation of these virulence factors is provided in Figure 5. Related to immune evasion, the protein Lysophospholipase L1 (COG2755), was detected exclusively in *G. piotii*, *G. pickettii* and GG14. Similarly, dTDP-4-dehydrorhamnose reductase was detected only in *G. vaginalis, G. piotii, G. pickettii,* and GG2. Moreover, allantoin utilization, mediated by allantoinase AIIB (K01466) and allantoin permease (K10975), was identified only in *G. leopoldii*, *G. swidsinskii*, GG13, and GG14. The vaginolysin (K11031), a thiol-activated cytolysin and member of the pore-forming toxin family, was detected in all isolates of *G. leopoldii*, *G. swidsinskii*, *G. greenwoodii*, GG9, GG10, and GG12-GG14, and in most of *G. vaginalis* (98.3%) and *G. pickettii* (82.61%), and more than half of GG2 (66.7%) isolates (Figure 5). The distribution was observed across strains isolated from humans with different health conditions. Proteins associated with mucin degradation, such as the α-/β-galactosidases LacZ (COG3250), GalA (COG3345), and endo-1,4-β-mannosidase (COG3934) were exclusive to *G. vaginalis* and GG2, while α-L-fucosidases were detected in *G. vaginalis*, GG2 and GG12. The α-mannosidase MngB protein was detected in all *G. vaginalis*, *G. pickettii*, *G. piotii*, GG2, and GG11 isolates, as well as in 66.7% of *G. swidsinskii* isolates. Sialidase NanH1 was detected in all *G. vaginalis*, *G. piotii*, *G. pickettii, G. greenwoodii,* GG2, GG9-GG11 and GG14 isolates, while NanH2 was present in 66.7% of *G. piotii* and 8.7% of *G. pickettii* (n=2/23) isolates, and in the unique GG11 isolate. The NanH3 was detected in all *G. piotii* and *G. pickettii*, and 18.9% of *G. vaginalis* isolates. Additionally, an uncharacterized exo-α-sialidase was detected in three *G. vaginalis* isolates and no sialidases were identified in *G.leopoldii, G. swidsinskii,* GG7, GG12 and GG13.

**Figure 5.**
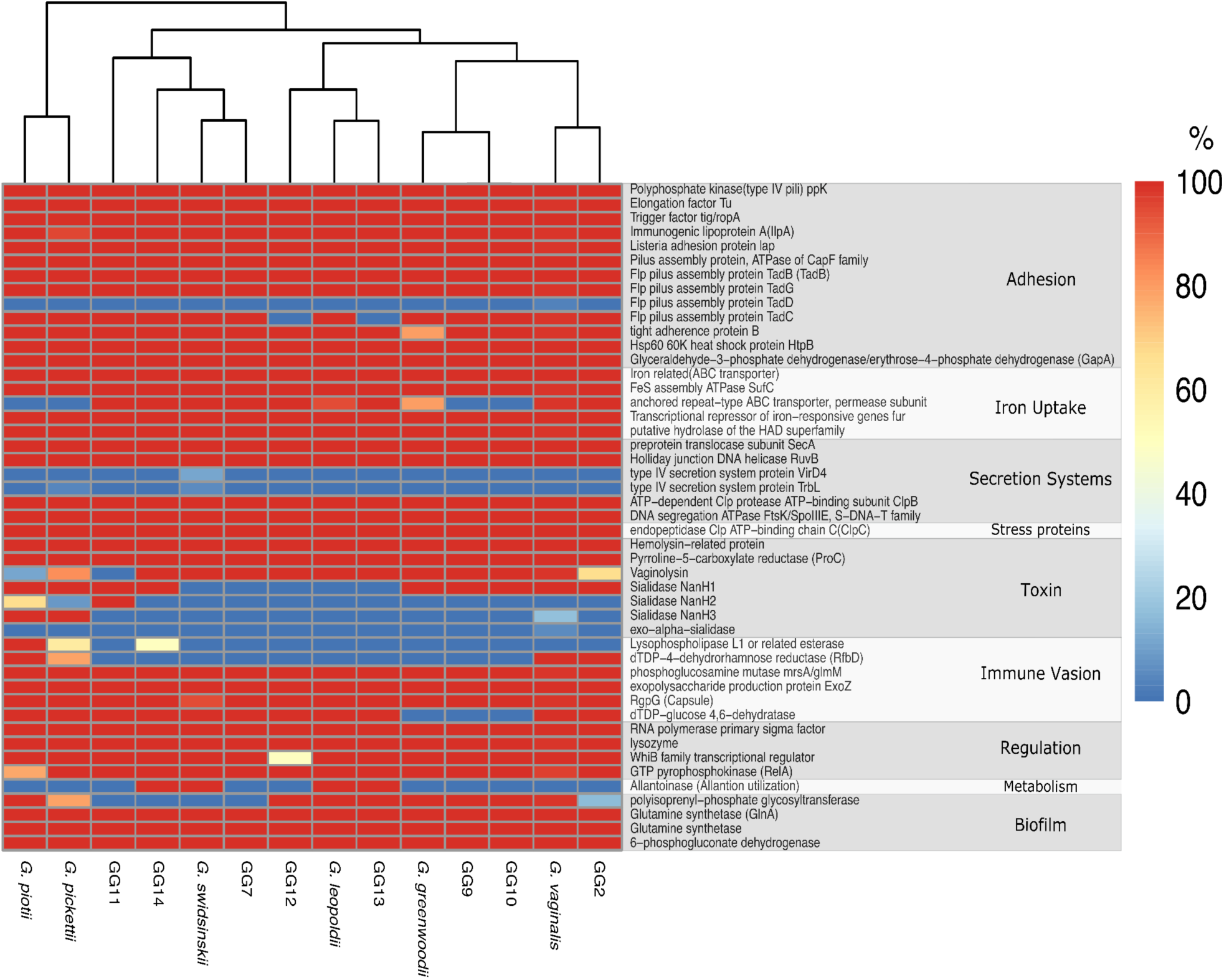
Heatmap showing the percentage of the most relevant VFs among *Gardnerella* genomes. The dendrogram represents the clustering of isolates based on Euclidean distance. The percentage of virulence genes is color-coded, with red (100%) indicating full presence and blue (0%) indicating absence. Intermediate prevalence is represented by shades of orange, yellow, and light blue.

Finally, four secondary metabolite biosynthetic gene clusters (BGCs) were identified across the 153 *Gardnerella* genomes (Table S7). All 18 *G. swidsinskii* isolates harboured a BGC encoding a ribosomally synthesised and post-translationally modified peptide product (RiPP), and approximately 83% (n=15/18) carried a BGC encoding a class II lanthipeptide. A BGC encoding a type III polyketide synthase was detected in all GG12 (n=2) and in most of *G. vaginalis* and *G. pickettii* isolates (93.1% and 82.6%, n=54/58 and n=19/23, respectively). Moreover, a BGC encoding a linear azol(in)-containing peptide was detected in *G. vaginalis* (3.4%, n=2/58) and *G. swidsinskii* (5.6%, n=1/18).

## Discussion

### Genomic Diversity and Species Delineation

The pathogenic potential of *Gardnerella* spp. has long been debated due to a lack of taxonomic resolution, previously grouping distinct species together. Advances in genomic analyses, including whole-genome sequencing and phylogenomics, now enabled a more detailed understanding of *Gardnerella*, revealing key differences in host-adaptations and pathogenic traits (21, 22, 26). This study presents the most comprehensive *Gardnerella* pangenome analysis to date, identifying a novel genomic species, GG15 (strains JCP7275 and N165), which had not been previously recognized (Figure 1). Our findings highlight that while ANI and dDDH values offer insight into genetic relatedness, they are insufficient for precise species delineation. Instead, core genome phylogeny provides a more robust framework for classification, while pangenome analysis highlights high intraspecies variability in genome size, coding sequence content, and metabolic capabilities.

The detection of the same *Gardnerella* clones in different biological samples from the same donor suggests that these species can colonize multiple anatomical sites within the urogenital tract and persist over time (54). The presence of *G. pickettii* clones in both asymptomatic women and those diagnosed with overactive bladder suggests a complex relationship between *Gardnerella* colonization and urogenital health. Similarly, the detection of specific *Gardnerella* species (*G. vaginalis, G. piotii, G. leopoldii* and *G. swidsinskii)* in vaginal swabs of women with BV (30) should not be interpreted as a direct causation, as other species, such as *Fannyhessea vaginae*, *Prevotella bivia*, are also associated with BV pathogenesis, emphasizing the polymicrobial nature of this condition (31).

Genome features analysis reveals high intraspecies variability in genome size and CDS, underscoring the genetic diversity within this genus (32). The overall *Gardnerella* pangenome metrics align with previous findings (4, 21), though minor discrepancies exist due to different pangenome computation tools, parameters, and the use of a larger genome collection, which tends to increase pangenome size while decreasing core genome size in species with open pangenomes (33).

The extensive accessory genome of *Gardnerella*, comprising 89.5% of its pangenome, reflects its high genetic diversity, a characteristic similar to *Bifidobacterium* species (22). This diversity is largely driven by metabolism-associated genes, which vary significantly among *Gardnerella* species.

### Metabolic Specialization and Host Adaptations

Distinct metabolic adaptations among *Gardnerella* species suggest niche specialization within the human urogenital tract. *G. vaginalis* and GG2 exhibit a high proportion of genes involved in carbohydrates transport and metabolism, particularly for glucuronate, pentose phosphate, and galactose degradation, key energy sources in the vaginal microbiome (34). The ability to metabolize pentose sugars and degrade galactose, a mucin component, may provide an advantage for colonization and persistence. Similarly, *G. piotii* and *G. pickettii* exhibit enrichment in carbohydrate metabolism genes, including those for keratan sulfate degradation, a glycosaminoglycan component of the urothelial glycocalyx (35). This capacity may facilitate bacterial persistence in the urinary tract, potentially linking these species to urogenital disorders such as overactive bladder.

Other features such as the lower abundance of translation, ribosomal structure and biogenesis associated proteins in *G. vaginalis* as well as the higher proportion of coenzyme transport and metabolism associated proteins in *G. leopoldii* and *G. swidsinskii.* These findings corroborate previous findings (23), and suggest that *G. vaginalis* may employ a different regulatory mechanism, while *G. leopoldii* and *G. swidsinskii* exhibit enhanced metabolic capabilities, potentially reflecting adaptations to the urogenital niche or host interactions. Moreover, the GG7, GG12, GG13, and GG14 appear to have a similar proportion of coenzyme transport and metabolism-associated proteins as *G. leopoldii* and *G. swidsinskii*. However, further genomes sequencing is required to validate these findings.

Beyond carbohydrate metabolism, differences in amino acid utilization may further drive species-specific adaptations. The identification of genes involved in L-tryptophan biosynthesis from indole in *G. leopoldii, G. swidsinskii,* GG7, and GG10 suggests a potential role in modulating vaginal health. Recent studies have shown that women with BV exhibit higher vaginal indole levels (36), indicating that metabolic shifts in the microbiota influence host-microbe interactions. If *Gardnerella* species with tryptophan biosynthesis genes contribute to indole metabolism, they may impact microbial balance, inflammation, or immune modulation in the vaginal ecosystem. Additionally, allantoin utilization genes detected in *G. leopoldii, G. swidsinskii,* GG13, and GG14 suggest an alternative carbon and nitrogen source that may provide a competitive advantage to obtain carbon and nitrogen from the urogenital environment (20, 37, 38)

### Virulence and Antibiotic Resistance

*Gardnerella* species harbour genes that may confer resistance to clinically relevant antibiotics, including those used to treat urogenital infections (37, 39, 40). This is particularly concerning because some antibiotic resistance genes (e.g. *lsa* (n=4), *mefa/msrD* (n=1)) were associated with mobile genetic elements (41), further suggesting the potential for horizontal gene transfer, necessitating closer monitoring of *Gardnerella* resistance mechanisms. Of note is the high prevalence of the *lsaC*-like gene, which may encode a protein able to confer resistance to clindamycin, a commonly prescribed antibiotic for BV (5, 37, 39, 40, 42).

The VF analysis was conducted using an *in house* database based on previous studies on *Gardnerella* (15, 37, 76) (Table S6). The content in VFs appears to be closely associated with *Gardnerella* speciation. The lysophospholipase L1 (COG2755) was detected exclusively in *G. piotii*, *G. pickettii* and GG14, warranting further investigation in the context of *Gardnerella* pathogenicity particularly because phospholipase, particularly phospholipase C, have been associated with BV (43). The presence of this VF suggests a possible mechanism for tissue invasion and colonization in the vaginal environment, potentially contributing to the disruption of the vaginal epithelium and the development of BV-associated symptoms (44). Additionally, the dTDP-4-dehydrorhamnose reductase (K23987) and 6-phosphogluconate dehydrogenase (K00033) were also found in *G. vaginalis*, *G. piotii*, *G. pickettii*, and GG2. The dTDP-4-dehydrorhamnose reductase, a key enzyme in the dTDP-L-rhamnose biosynthesis pathway, has been shown to impact either the viability or virulence of numerous bacterial species (45). Similarly, 6-phosphogluconate dehydrogenase (K00033), a surface-located immunogenic lectin protein, can function as an adhesin and has demonstrated significance in bacterial pathogenesis (46, 47).

Previously identified as an accessory gene (17), vaginolysin was widely distributed across *Gardnerella* species in this study, including strains from humans with diverse health conditions. Vaginolysin, a human-specific cytotoxin, binds to the complement regulatory protein CD59, leading to pore formation in host cell membranes (48). Furthermore, its cytolytic activity releases intracellular contents, potentially providing nutrients that promote the growth of *Gardnerella* and other BV-associated microorganisms (49). However, some strains lacking vaginolysin likely compensate for its absence through alternative mechanisms, suggesting functional redundancy and adaptability within the genus.

Proteins associated with mucin degradation, particularly NanH2 and NanH3, which degrade sialic acids on epithelial cells, were predominantly found in *G. vaginalis, G. piotii,* and *G. pickettii* (15). NanH2 and NanH3, have been previously recognized as key contributors to sialidase activity, directly involved in breaking down sialic acids (15). This enzymatic activity may lead to glycosaminoglycan layer depletion, compromising the vaginal mucosal barrier and potentially increasing the risk of bacterial adherence and infection (15, 50). However, these proteins have also been identified in healthy women, suggesting their presence alone may not be sufficient to drive pathogenesis (15). Although previously BV-associated, NanH1 appears to exhibit basal sialidase activity (15, 50), likely contributing more to general metabolic processes rather than active mucin degradation. Additionally, the uncharacterized exo-α-sialidase identified in this study, likely corresponding to the recently identified NanH4, may be inactive, as it is present in *G. vaginalis* ATCC 49145, a strain that does not exhibit sialidase activity (17). However, functional sialidase activity assays are necessary to confirm this prediction and determine its potential role in *Gardnerella* physiology. These results suggest that sialidases may provide a competitive advantage in the urogenital environment by enabling *Gardnerella* to utilize a broader range of energy sources.

Moreover, the utilisation of allantoin as an intermediate metabolite of purine metabolism is a trait shared by some bacterial species, which can contribute to their virulence (38, 51), by playing a role in iron acquisition, a crucial element for bacterial growth and survival (38).

### Secondary Metabolite Biosynthetic Gene Clusters

The identification of BGCs across *Gardnerella* genomes suggests that these species may produce bioactive compounds with potential ecological or pathogenic significance. All *G. swidsinskii* isolates harbored a BGC encoding a ribosomally synthesized and post-translationally modified peptide product (RiPP), while most *G. vaginalis* and *G. pickettii* isolates carried a BGC for a type III polyketide synthase. Additionally, a linear azol(in)-containing peptide BGC was detected in a small subset of *G. vaginalis* and *G. swidsinskii* isolates. The role of these secondary metabolites in *Gardnerella* physiology remains unclear, but their presence suggests potential antimicrobial properties or involvement in host interactions.

### Conclusions

This study provides the most detailed genomic characterization of *Gardnerella* genus, uncovering extensive genetic diversity, metabolic versatility, and species-specific adaptations. The identification of a previously unrecognized genomic species (GG15) further refines our understanding of *Gardnerella* taxonomy. Approximately 40% of the genes in the *Gardnerella* pangenome are associated with known COG functions, while the genus’s extensive accessory genome—comprising 90% of the total pangenome—highlights its significant metabolic versatility, which may contribute to its dual roles in both health and disease. NMDS analysis further differentiated *G. vaginalis* and GG2 from other *Gardnerella* species, highlighting their unique genetic traits, particularly the higher proportion of proteins involved in carbohydrate transport and metabolism. This suggests that these species may have specialized metabolic pathways that enable them to exploit specific carbohydrate sources in the urogenital environment, potentially influencing their survival and colonization, which could play a role in their persistence and pathogenicity. Understanding the metabolic profiles of these species could provide insights into their ecological niches and offer new avenues for targeting species-specific therapeutic strategies. Our analysis identified species-specific virulence factors, such as allantoin utilization in *G. leopoldii* and *G. swidsinskii*, which may enable these species to thrive in the urogenital tract. Importantly, they have the potential to offer novel therapeutic or diagnostic targets for conditions like bacterial vaginosis.The presence of antimicrobial resistance genes (ARGs), such as *lsa*(C), raises concerns about the effectiveness of current treatments and the risk of resistance spread. Moreover, the detection of identical *Gardnerella* clones in both urine and vaginal samples suggests the possibility of persistent colonization across different urogenital sites. This may indicate the capacity for *Gardnerella* species to establish stable, long-term associations within the host microbiota, which could influence the dynamics of *Gardnerella* in both health and disease. However, there is currently no direct evidence linking these specific clones to the onset or recurrence of bacterial vaginosis (BV), warranting further investigation into their potential role in disease persistence or recurrent infections.

The lack of comprehensive host health data impacts the interpretation of pangenome findings. Host health information is crucial for understanding the intricate interactions between *Gardnerella* and the urogenital tract. Recent research has shown that Gardn*erella* exposures can alter bladder gene expression, affecting inflammation, immunity, and epithelial turnover (52, 53). These host responses may determine whether *Gardnerella* colonization leads to symptomatic conditions or persists in a commensal state.

Future research should prioritize the complex ecological dynamics between *Gardnerella* species and other microorganisms within the urogenital microbiome. A deeper understanding of the pathogenic mechanisms of different *Gardnerella* species across various urogenital conditions will be essential for the development of targeted, species-specific diagnostic and therapeutic strategies. Additionally, sequencing more *Gardnerella* genomes, particularly from under-studied species, and investigating host-pathogen interactions at the molecular level. Furthermore, exploring the role of *Gardnerella* in increasing susceptibility to other urogenital pathogens will provide deeper insights into its clinical significance.

## Materials and Methods

### Bacterial Collection

Thirty-three *Gardnerella* strains were isolated from urine and vaginal swabs of eight asymptomatic women (one postmenopausal) and three women diagnosed with OAB (one postmenopausal). Participants provided midstream urine (0.1 mL) and vaginal swabs (54). Samples were processed as previously described, and preliminary identification was based on *cpn60* sequencing (54), resulting in *G. vaginalis* (n=12), *G. pickettii* (n=7), *G. greenwoodii* (n=1), *G. leopoldii* (n=7), *G. swidsinskii* (n=6). Genomic DNA was extracted as previously described (2), and the strains were subsequently subjected to whole-genome sequencing.

### DNA Extraction and Whole-Genome Sequencing

Isolates were grown on 30 ml of Brain Heart Infusion broth (Frilabo, Portugal) supplemented with 1% glucose, 1% starch, 0.5% yeast extract and 1% gelatin, and 5% sheep blood (Thermo Fisher Scientific, USA). Cultures were incubated at 37 °C in 5% CO_2_ for 48 h. Genomic DNA was extracted using the Wizard® Genomic DNA Purification Kit (Promega, USA) with the following modifications: 30 ml of culture was centrifuged for 10 min at 5000 g to *pellet* cells, washed with PBS, and treated with lysozyme for 20 min at 37°C, followed by addition of 10% SDS and proteinase K under the same conditions. DNA concentration was measured using the Qubit™ dsDNA High Sensitivity Assay Kit (Invitrogen™, Thermo Fisher Scientific, USA) and integrity confirmed on a 0.8% agarose gel.

Whole-genome sequencing was performed by Eurofins Genomics (Constance, Germany) using the NEBNext Ultra II FS DNA library prep kit (England Biolabs, San Diego, CA, USA) and sequencing on a NovaSeq 6000 instrument (Illumina) producing 2x150-bp paired-end reads. Reads were trimmed by Trim Galore 0.6.6 (https://www.bioinformatics.babraham.ac.uk/projects/trim_galore/) in Bacterial and Viral Bioinformatics Resource Center (https://www.bv-brc.org/)(55) and quality checked using FastQC 0.11.9 (56). *De novo* assembly was performed using Unicycler v0.4.8 (57) and assembly quality assessed by QUAST v5.0.2 (58). Genome annotation was generated through the NCBI Prokaryotic Genome Annotation Pipeline (59).

### *Gardnerella* Genomes Dataset

Genome sequences of *Gardnerella* strains were downloaded from the RefSeq NCBI public database (as of May of 2023). After performing quality control to remove low-quality, incomplete assemblies, and duplicated genomes, a total of 120 high-quality genomes were retained for downstream analysis (60). These genomes were annotated with Prokka v1.13 to ensure comparable quality standards (61). The 33 genomes sequenced in this study were added to this collection, resulting in a final dataset of 153 genomes.

### Pangenome Analysis

The pangenome of the *Gardnerella* genus was generated by identifying homologous genes through Markov clustering with an inflation parameter of 6 in anvi’o v7.1 (Distance: Euclidean; Linkage: Ward) (62). Predicted protein sequences were annotated against the Clusters of Orthologous Groups of proteins (COG) and Kyoto Encyclopedia of Genes and Genomes (KEGG) databases (63, 64). For each genome, the percentage of proteins in each COG category was calculated, and the distribution of the proportional abundance of each COG category was compared between *Gardnerella* species using the Kruskal-Wallis test. Multiple comparisons between two independent *Gardnerella* species/GG were performed using the unpaired two-sample Wilcoxon test. The structural patterns of COG categories and *Gardnerella* species/GG were visualized using Non-metric Multidimensional Scaling (NMDS) based on Bray-Curtis dissimilarity matrices. Similarity Percentages (SIMPER) were calculated with 999 permutations to identify which COG categories were responsible for differences between *Gardnerella* spp. (65). Functional enrichment analysis was performed to infer the enrichment scores of metabolic modules and pathways associated within specific genomes (66). A function or metabolic module was enriched in each group if the adjusted q-value was less than 0.05 and present in at least 50% of the *Gardnerella* strains.

Pangenome analysis of particular *Gardnerella* species (minimum of 10 genomes), namely *G. vaginalis*, *G. leopoldii*, *G. swidsinskii* and *G. pickettii*, was generated with Roary (67). Single Nucleotide Polymorphisms (SNPs) were extracted from the core genome alignment of each *Gardnerella* spp. using SNP-sites, and the SNP-alignment was used to generate a phylogenetic tree with RAxML using the general time-reversible model and gamma correction with bootstrap support of 1000 replicates (68, 69). The resulting core genome SNP tree was combined with a presence/ absence matrix of core and accessory genes in each pangenome. Core genome alignments were also used to calculate pairwise SNP distance matrices using snp-dists (https://github.com/tseemann/snp-dists). The α value of Heap’s Law was calculated for each *Gardnerella* spp. based to assess genomic plasticity, based on pangenome results from Roary.

### Taxonomic and Core Genome Analysis

Average Nucleotide Identity (ANI) and digital DNA-DNA hybridization (dDDH) values were calculated between the 153 *Gardnerella* genomes using fastANI and the Type Strain Genome Server platform, respectively (70, 71). Network analysis was conducted usingCytoscape v3.9.1 with the recommended thresholds for species delineation (70% for dDDH and 96% for ANI) (27–29). A core genome phylogeny was computed by aligning the concatenated amino acid sequences of 92 single-copy core genes using anvi’o (62). FastTree and iTOL v5 were used to compute and visualize the maximum-likelihood core genome tree, respectively (72, 73).

### Genome Mining of Antibiotic Resistance, Virulence and Secondary Metabolites

Antibiotic resistance genes were detected using the Comprehensive Antimicrobial Resistance Database (CARD) and the AMRFinder v3.10.45, applying a minimum identity threshold of 90% and 100% coverage (74, 75). A database of virulence factors previously described in *Gardnerella* was used for genome mining against the *Gardnerella* genus pangenome (15, 37, 76). The proportion of virulence factors represented in each *Gardnerella* species/GG was visualized as a heatmap, generated using the heatmap library with Euclidean distances and complete clustering (77). Secondary metabolite biosynthetic gene clusters were identified using antiSMASH v5 (78).

## Author Contributions

PT designed the study, performed the genomic and pangenome analysis, and wrote part of the original draft, MS performed the isolates isolation, genomic DNA extraction, annotation, and species identification, and wrote part of the original draft of the manuscript. FG and TGR contributed to study design, reviewing the manuscript and supervision. LP contributed to reviewing the manuscript, project design, administration, and funding.

## Ethical Approval

Approval of the study was obtained from the Faculty of Pharmacy (University of Porto, Porto, Portugal) Ethics Committee. Procedures performed in the study were all in accordance with the ethical standards of the institutional and national research committee, with the 1964 Helsinki Declaration, and its later amendments. All individual participants included in the study had given written informed consent.

## Supplementary Material

**Fig. S1 – S6** and **Table S1 - S7**: Supplementary information regarding *Gardnerella* genome data used in the current study.

**Figure S1.**
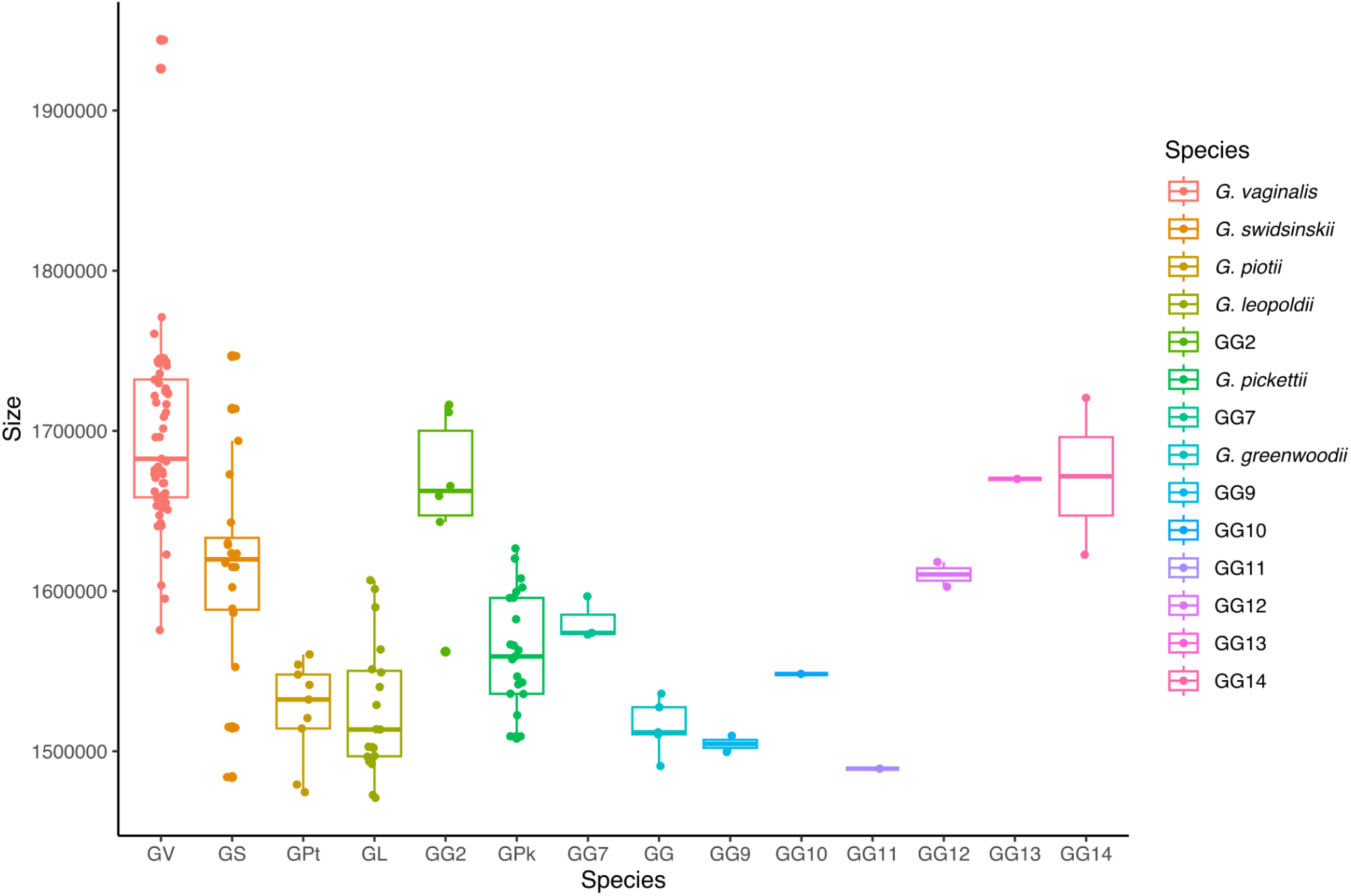
Average genome size across *Gardnerella* species and genomic species, ranging from 1.50 ± 0.02 Mbp (*Gardnerella* genomic species 9 - GG9) to 1.69 ± 0.06 Mbp (*G. vaginalis*) (p value < 2.2x10^-16^), with an overall average of 1.61 ± 0.09 Mbp.

**Figure S2.**
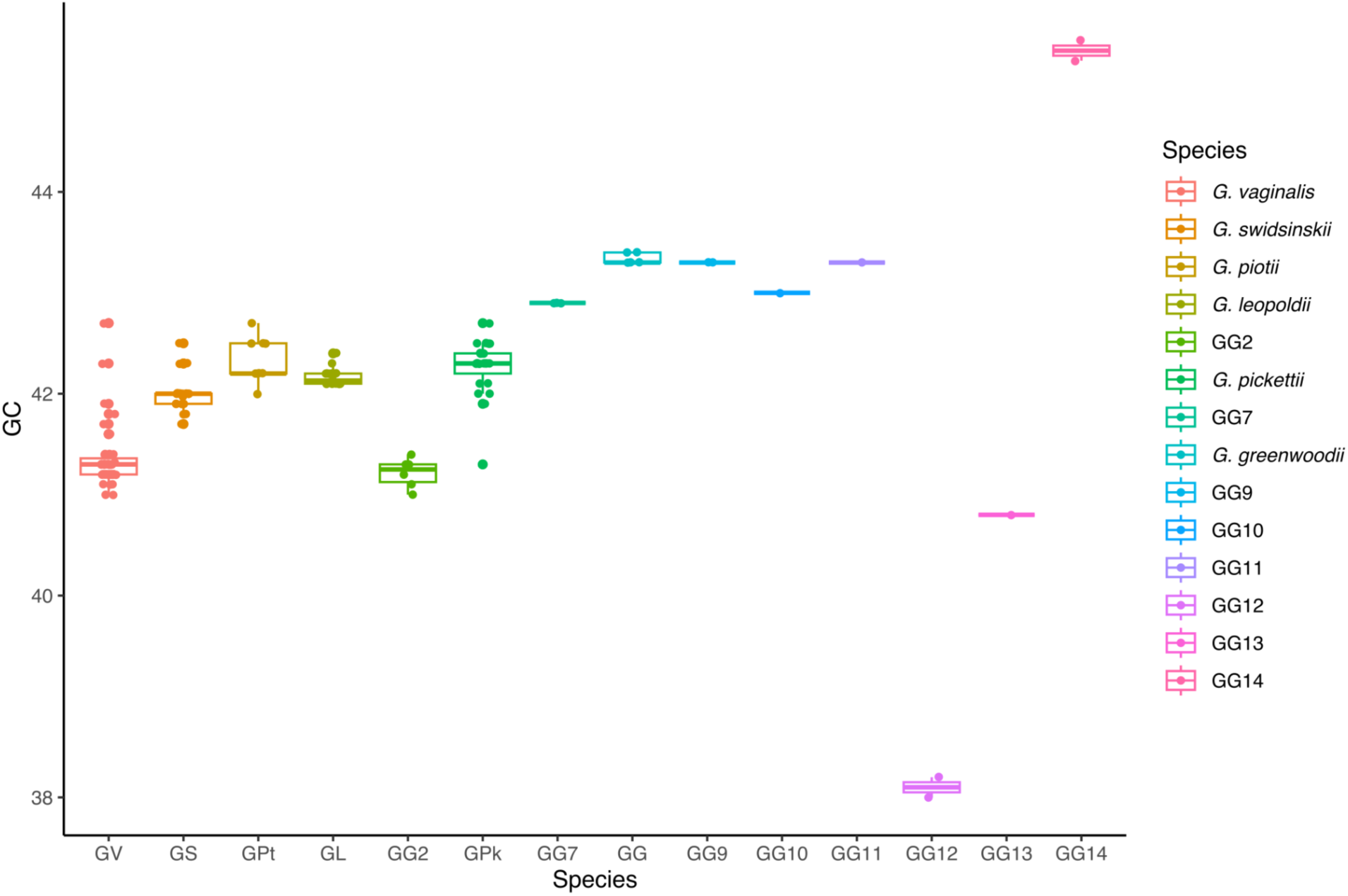
Guanine-Cytosine (GC) content across *Gardnerella* species and genomic species. *Gardnerella* genomic species 12 (GG12) genomes exhibit the lowest GC content at 38.1 ± 0.1%, while *Gardnerella* genomic species 14 (GG14) genomes display the highest at 45.4 ± 0.1% (p value < 2.2x10^-16^).

**Figure S3.**
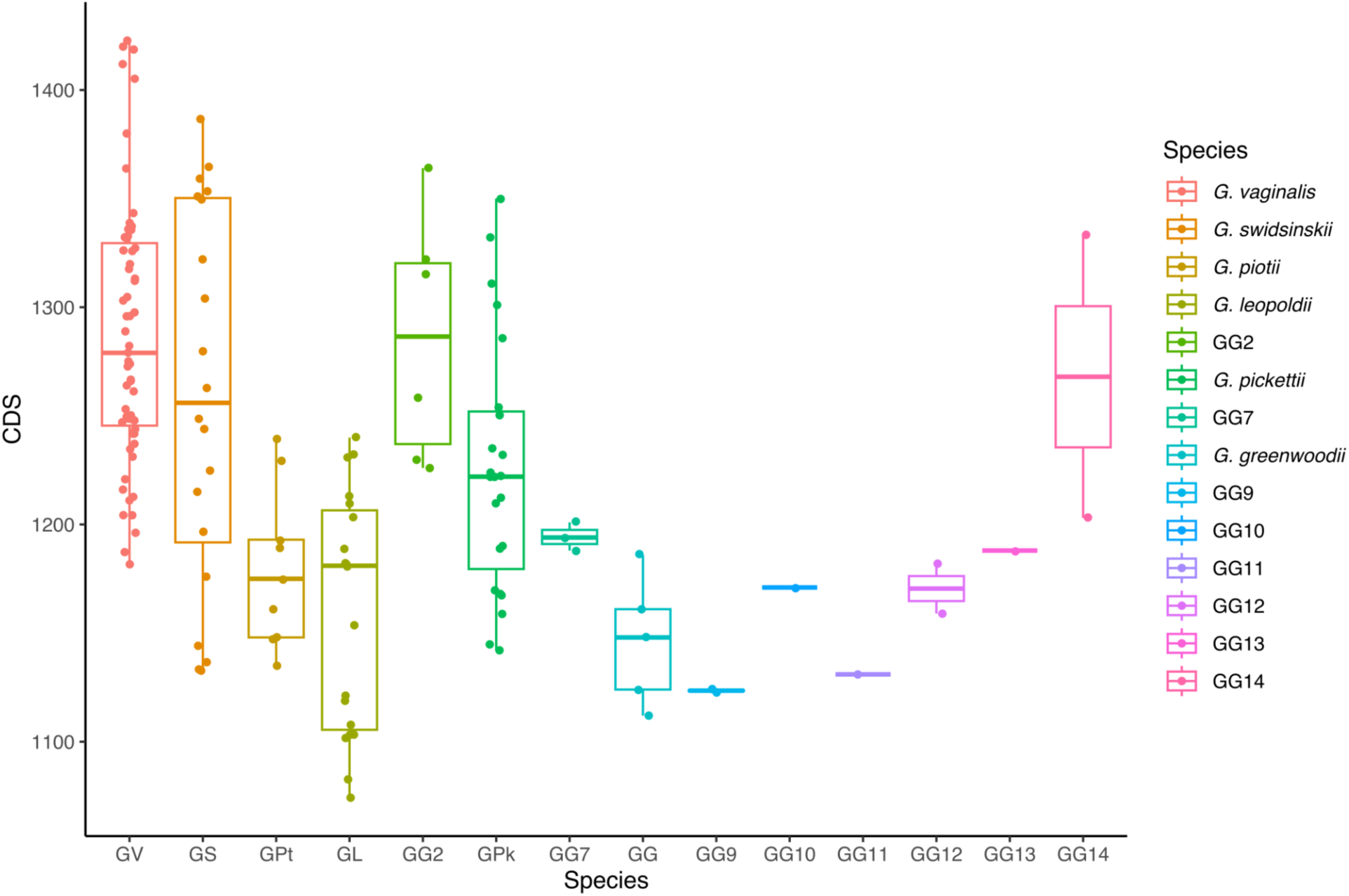
Number of coding sequences (CDS) across *Gardnerella* species and genomic species, ranging from 1124 ± 0.7 (*Gardnerella* genomic species 9 - GG9) to 1287 ± 61 (*G. vaginalis*) (p value < 6.8x10^-10^).

**Figure S4.**
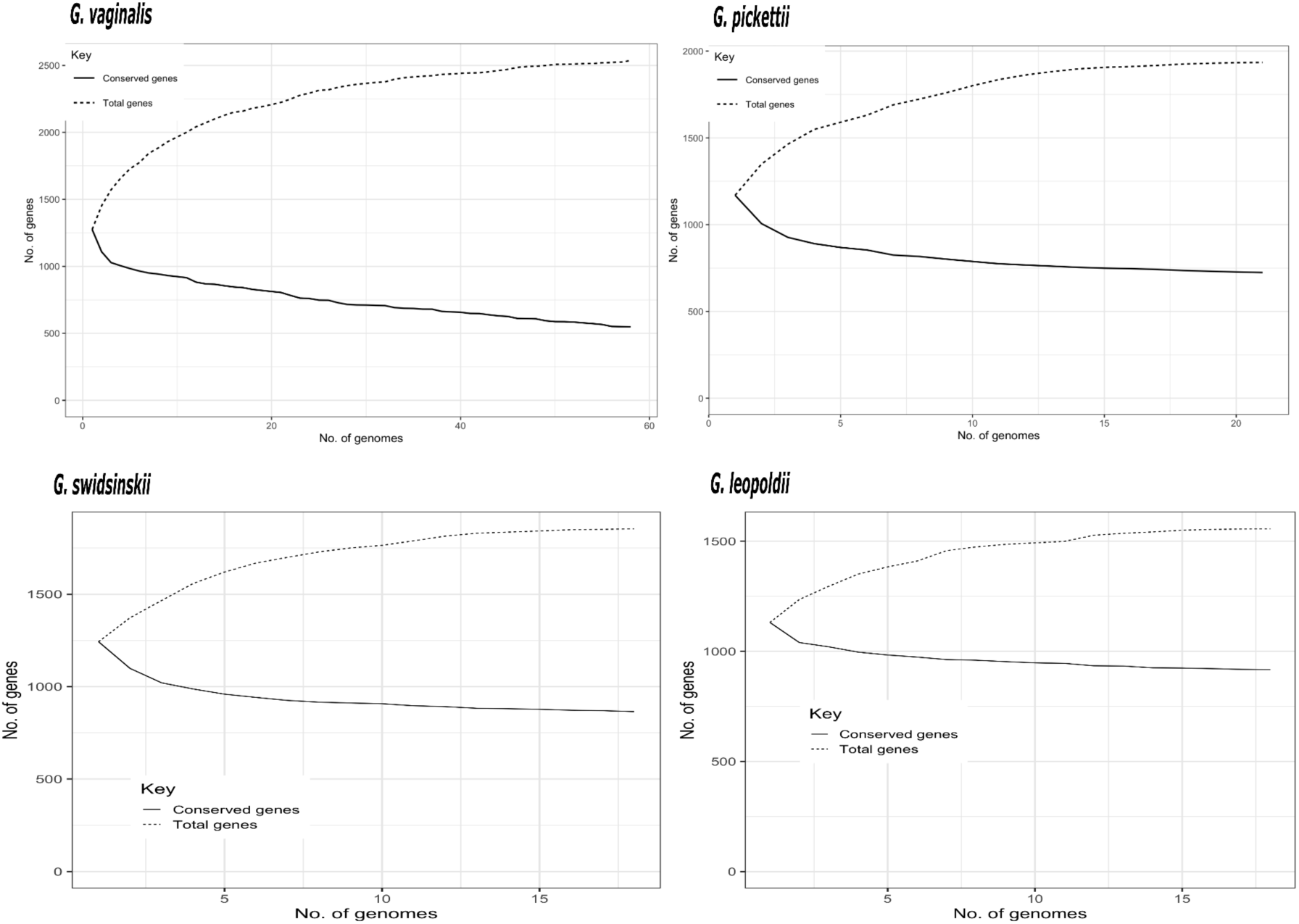
Pangenome analyses for *G. vaginalis, G. leopoldii, G. swidsinskii,* and *G. pickettii* revealed that all these species exhibit open pangenomes.

**Figure S5.**
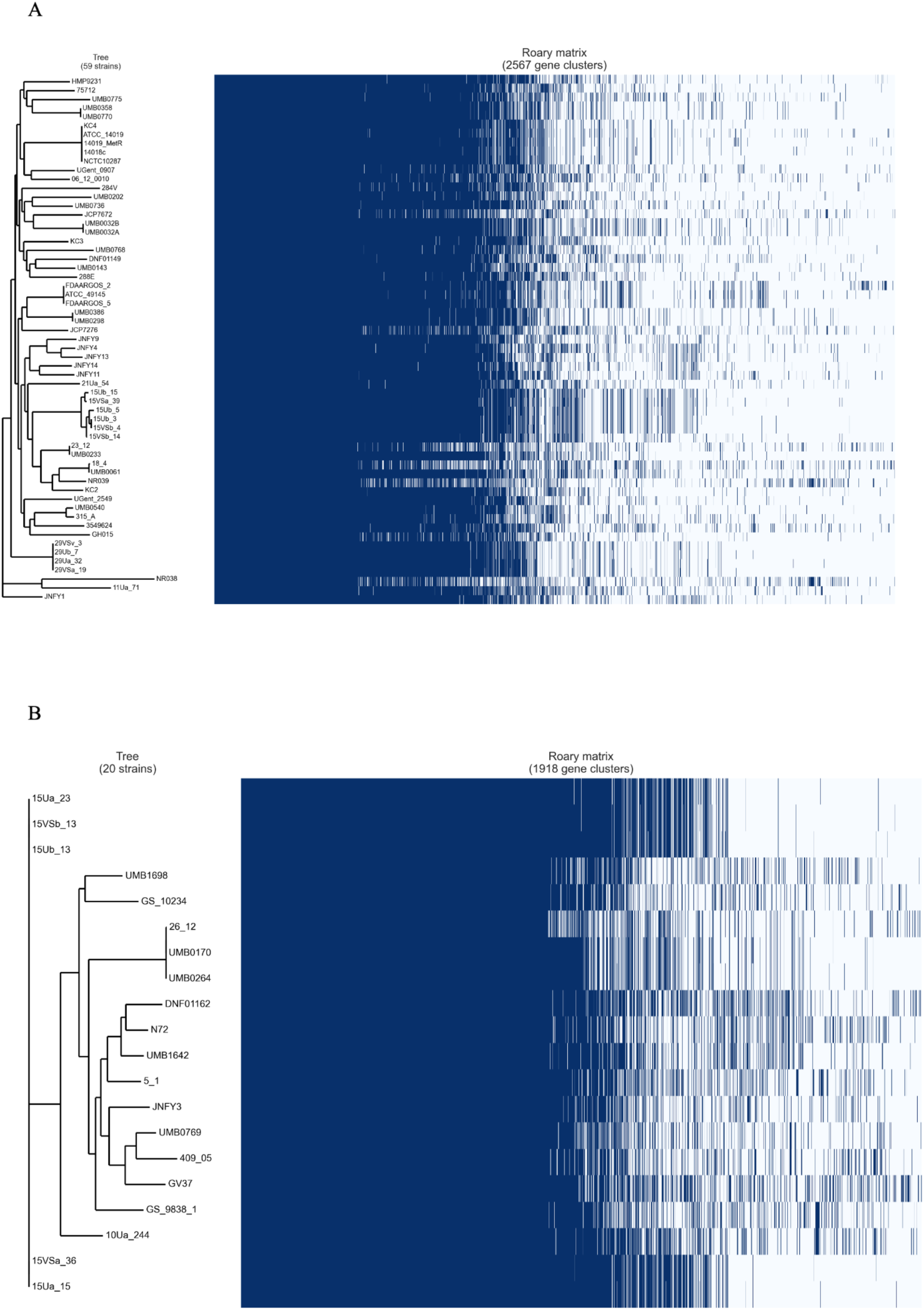
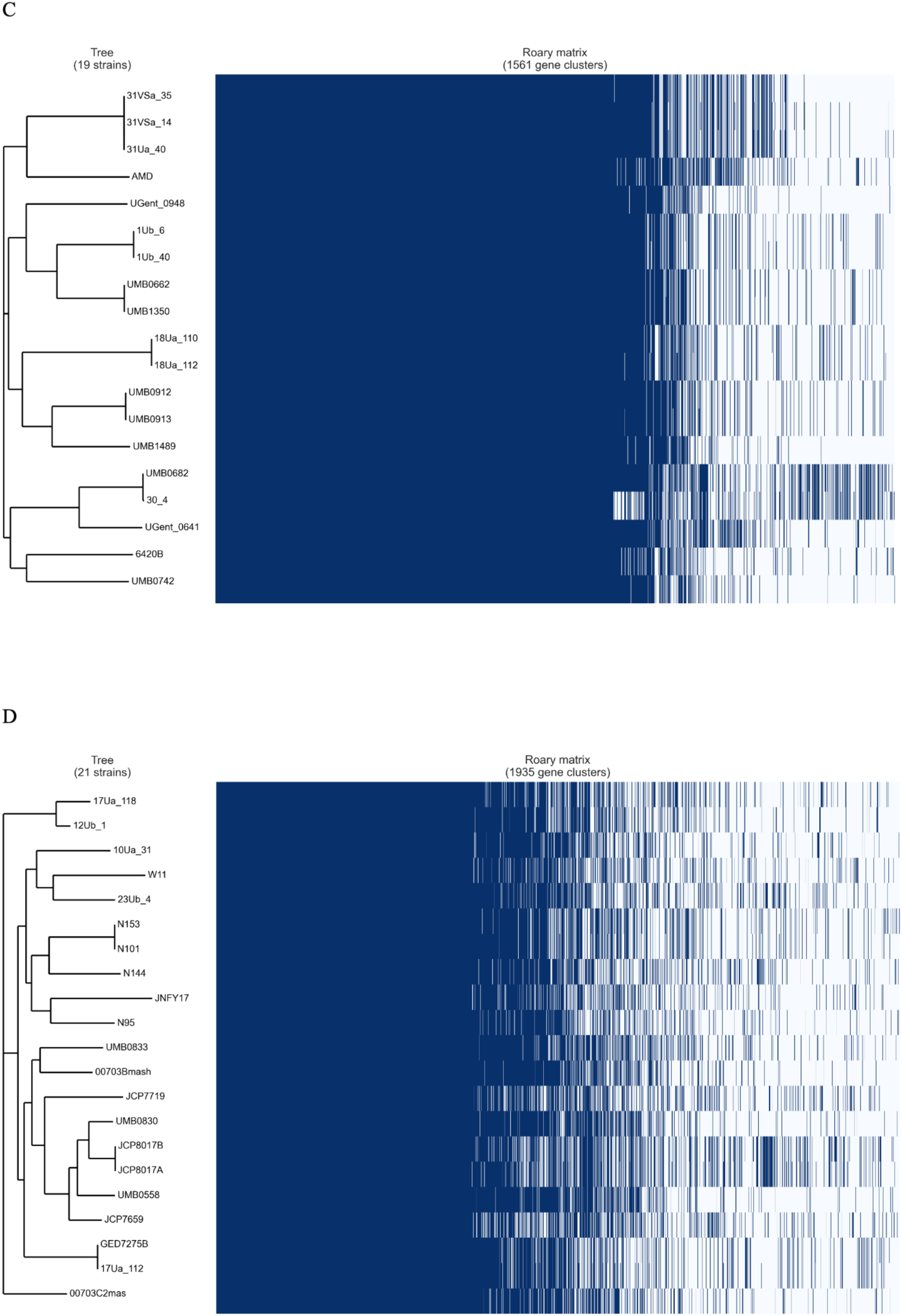
Core phylogeny and SNP matrix analysis, we identified identical clonal strains present in distinct biological samples from the same donor, specifically urine and vaginal swabs, across *G. vaginalis* (A)*, G. swidsinskii* (B)*, G. leopoldii* (C) and *G. pickettii* (D), species.

**Figure S6.**
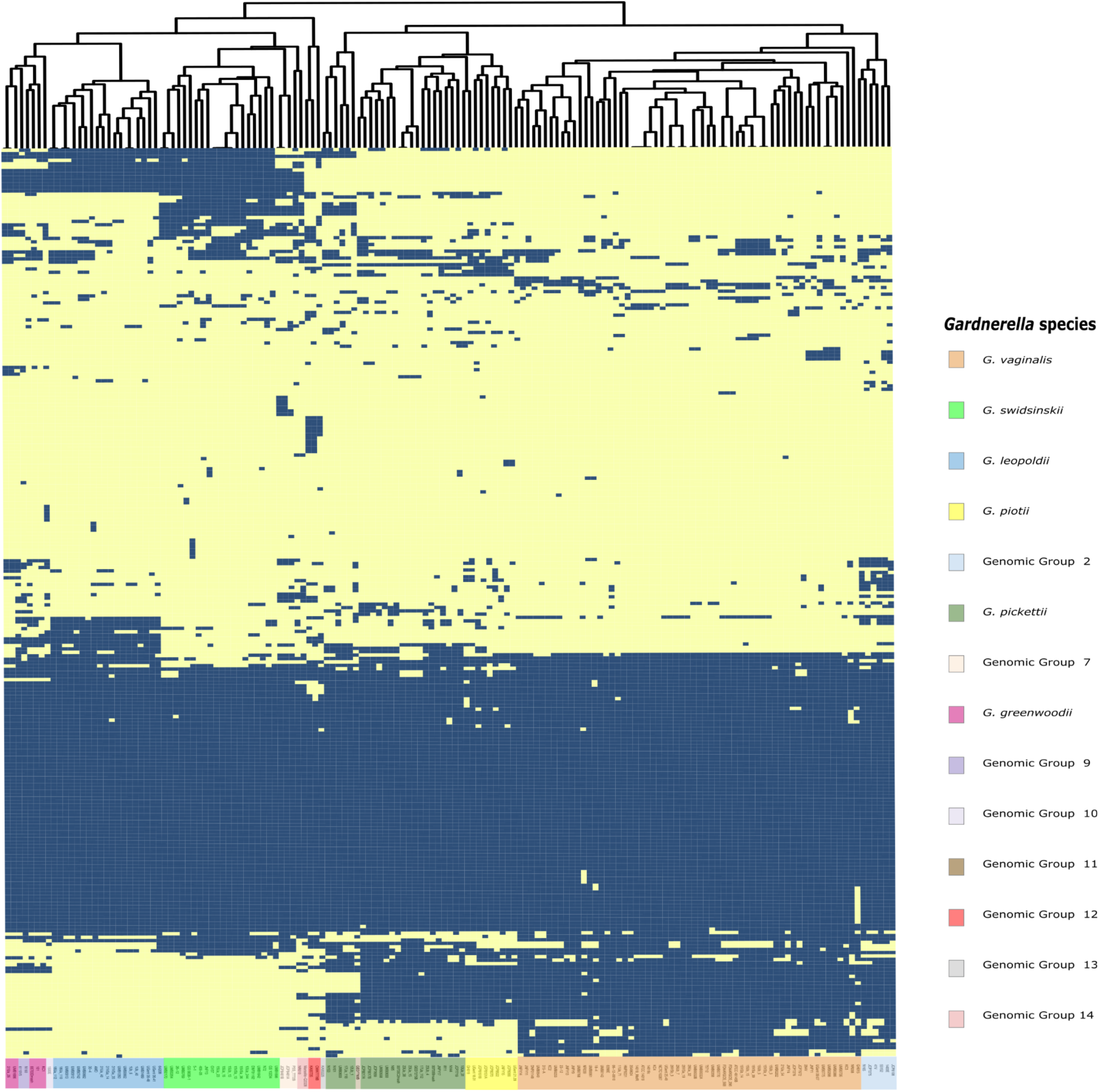
Heatmap representing presence/absence of putative VFs among *Gardnerella* genomes. The dendrogram results from clustering of isolates based on Euclidean distance. Presence or absence of virulence genes is represented as blue and yellow, respectively.

**Table S1:**
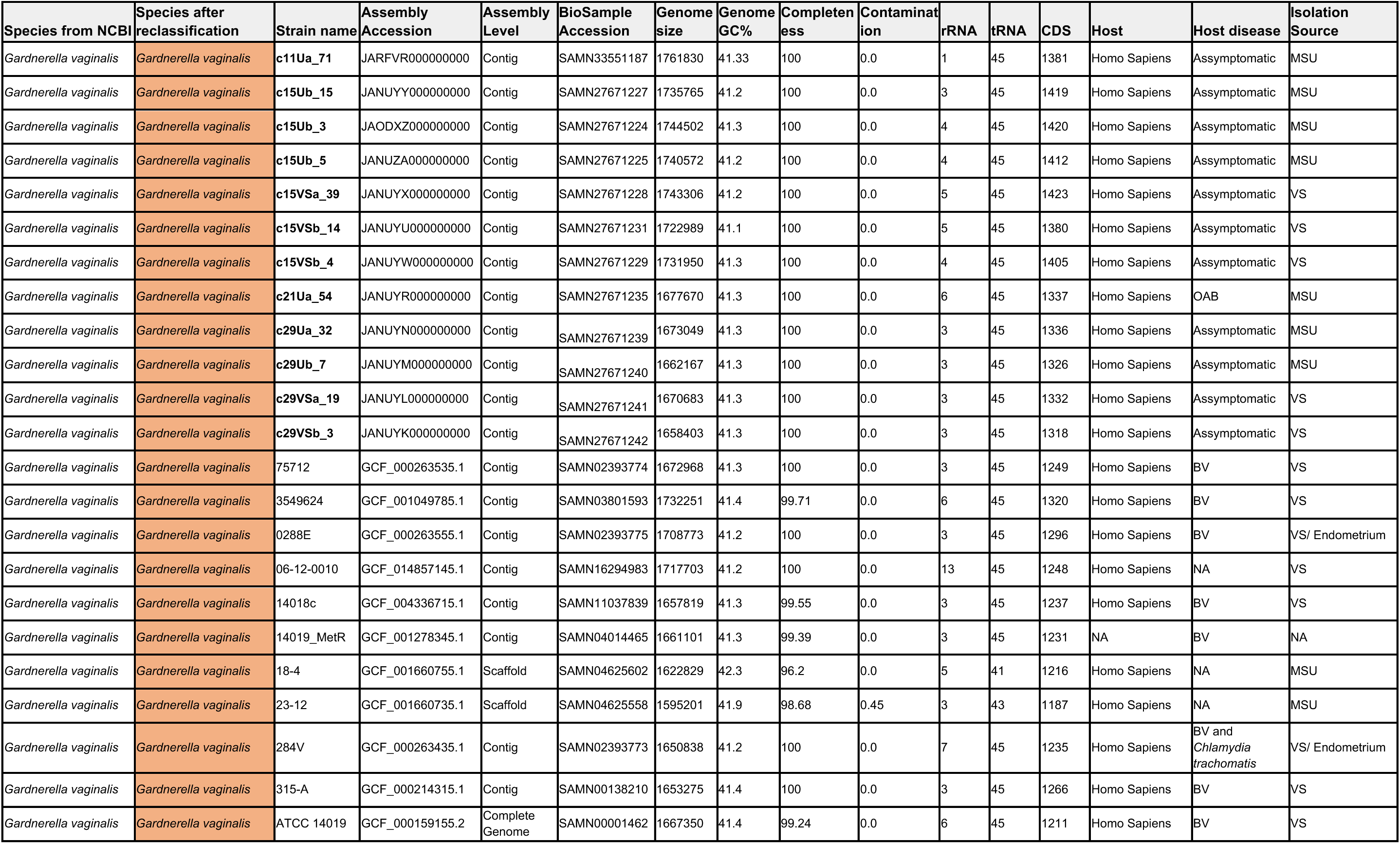
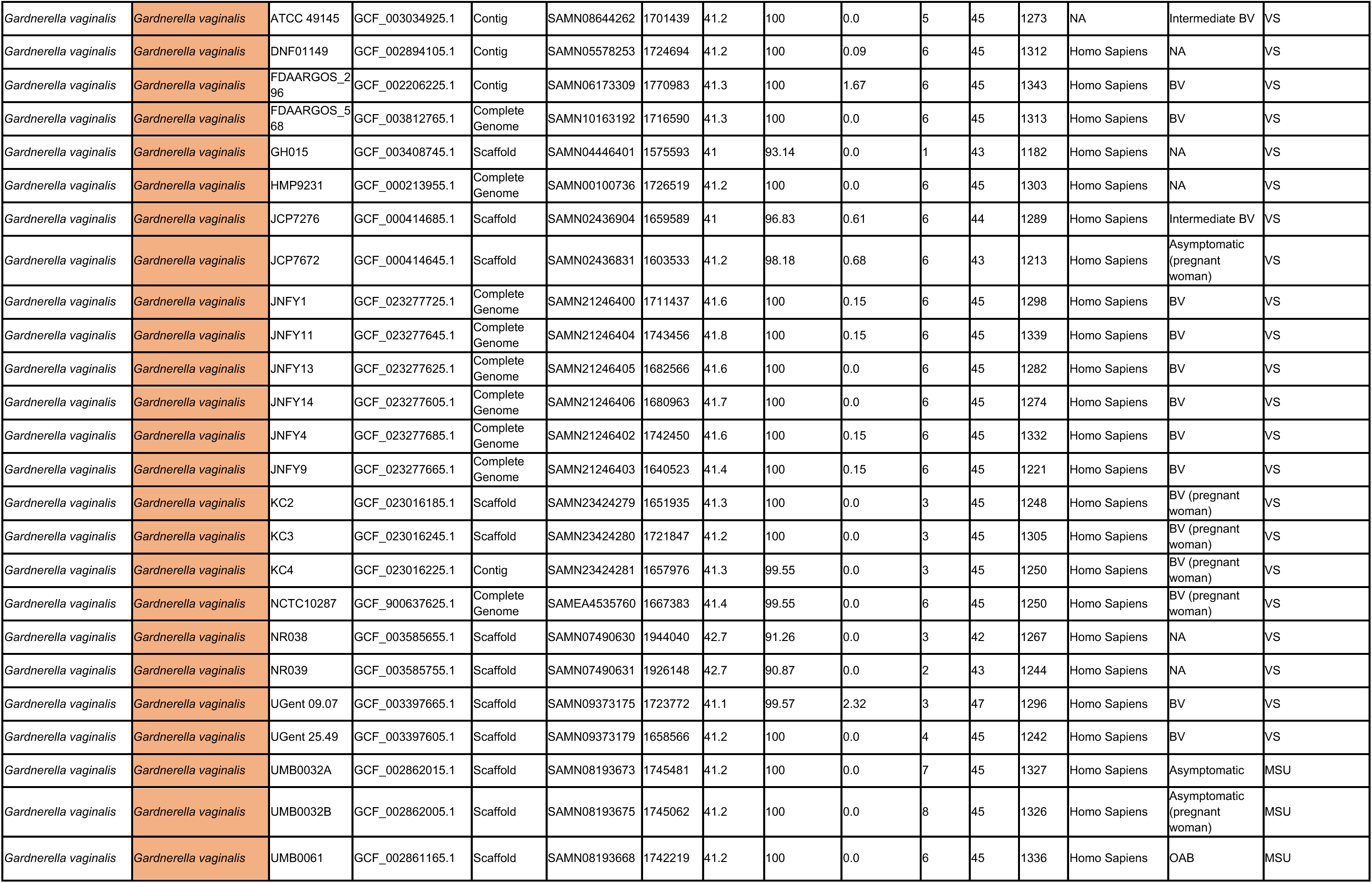
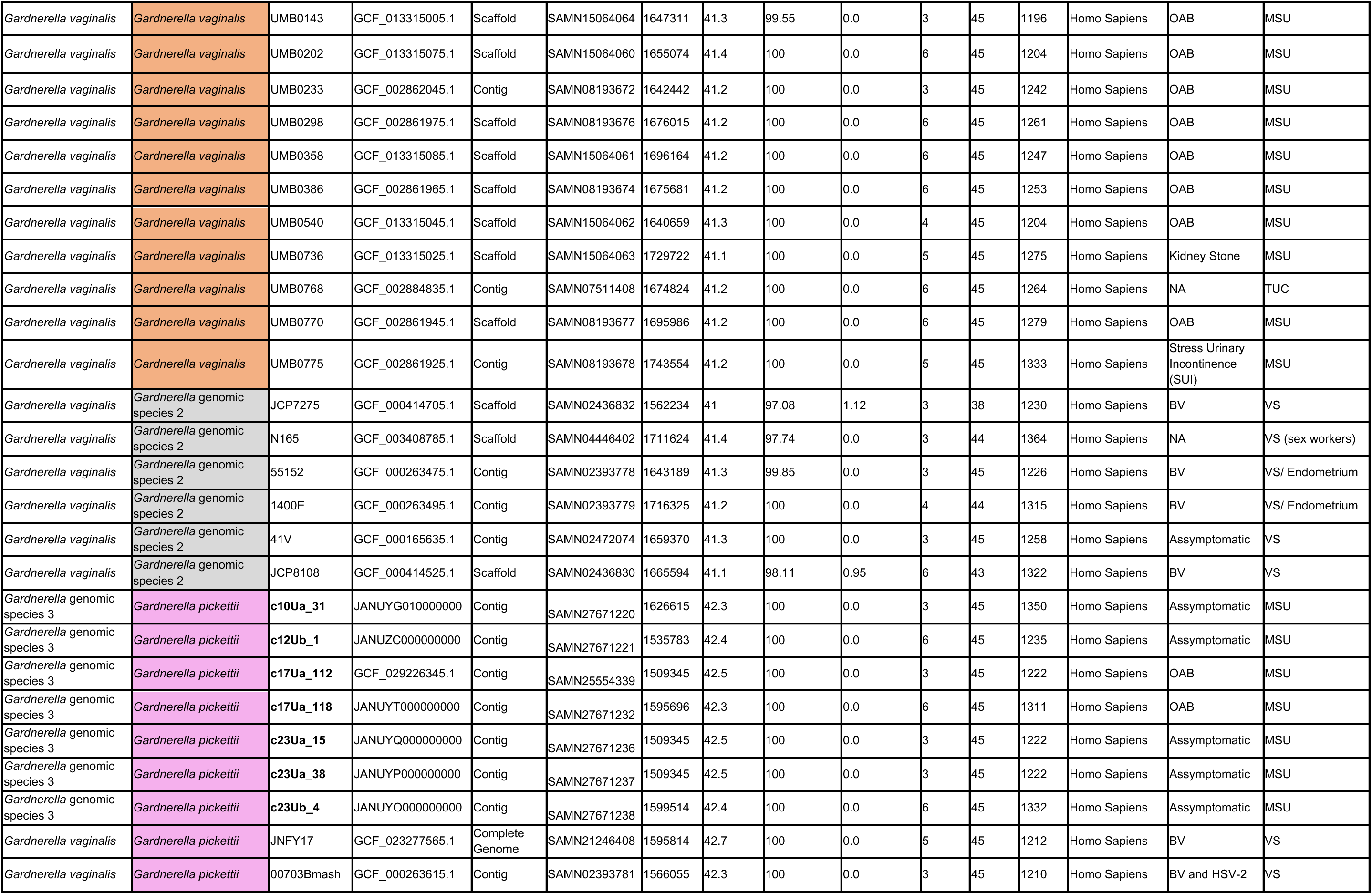
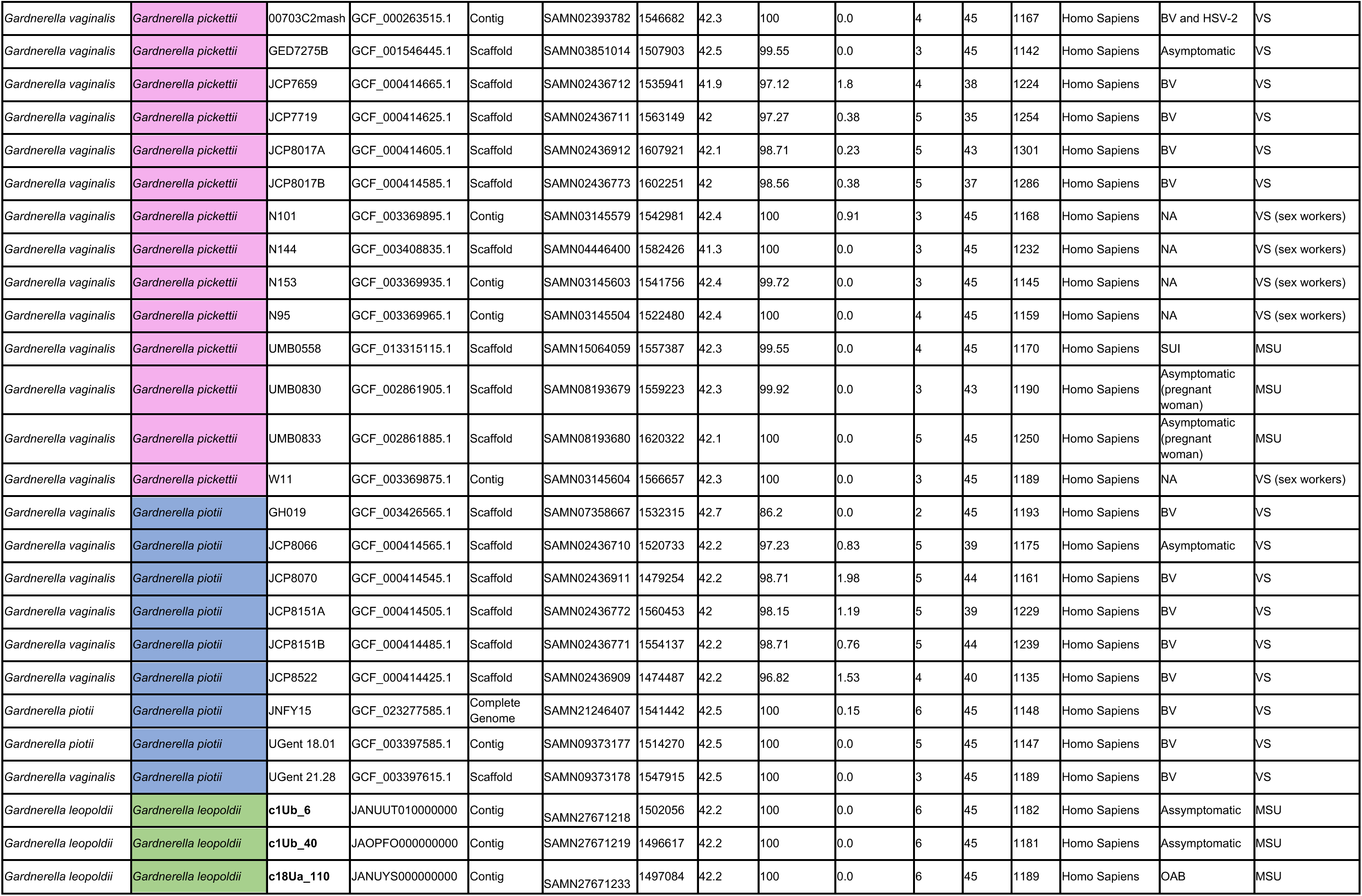
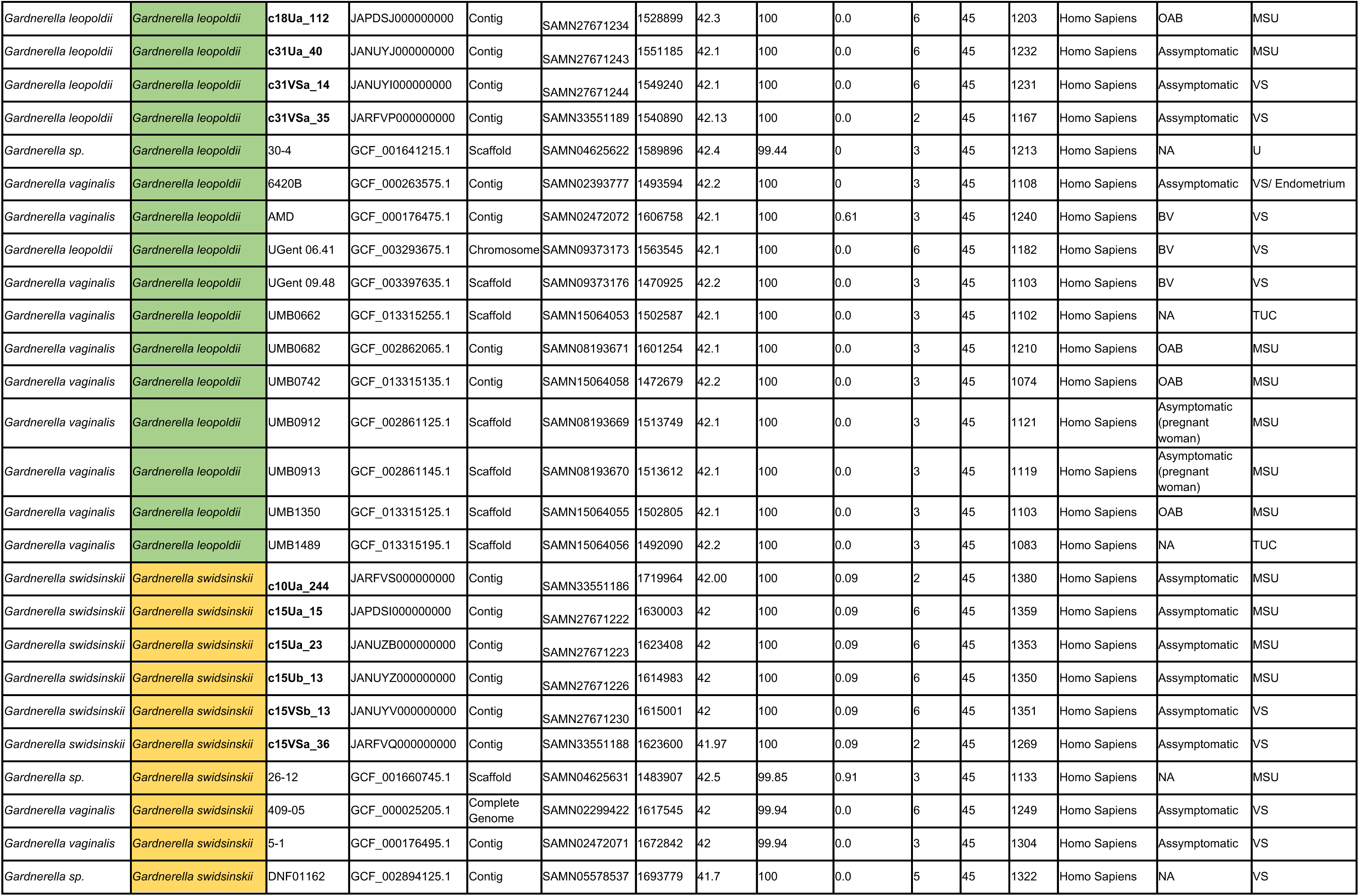
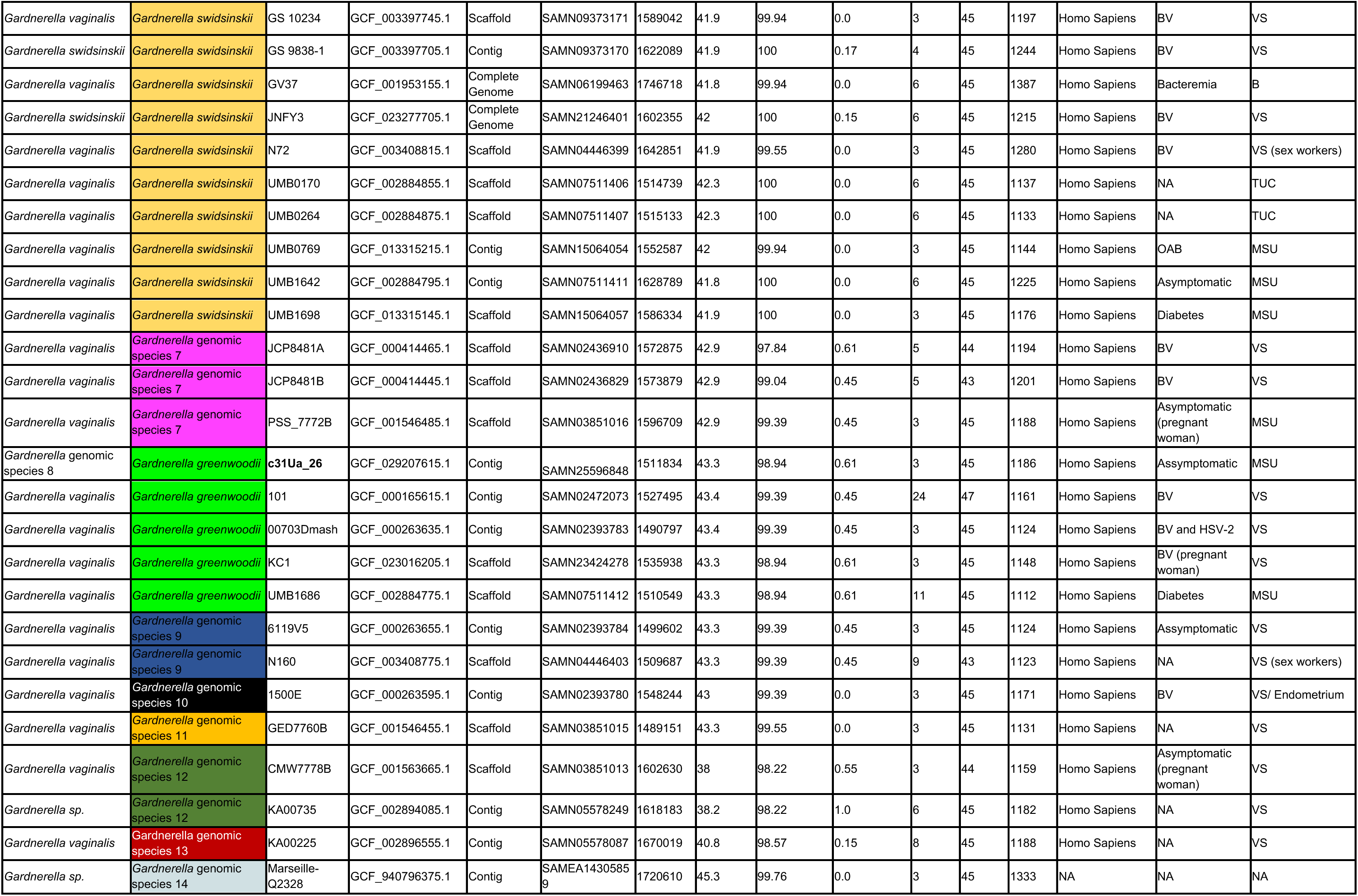
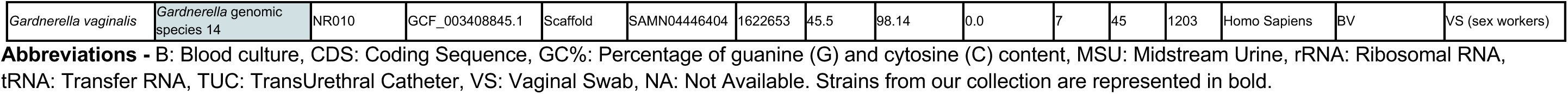
Genomic and metadata overview of reclassified *Gardnerella* strains: *Gardnerella* strains from our collection (boldtype) and genomes retrieved from NCBI (available in May 2023).

**Table S2:** Comprehensive analysis of the *Gardnerella* pangenome: core, soft core, dispensable, and strain-specific genes.

|  | Gene clusters | Gene Calls | % |
| --- | --- | --- | --- |
| **Core** | 505 | 77421 | 10,5 |
| **Soft Core** | 316 | 47316 | 6,6 |
| **Singletons** | 1419 | 1437 | 29,4 |
| **Dispensable** | 2584 | 65109 | 53,6 |
| **Accessory** | 4319 | 113862 | 89,5 |
| **Single copy genes in Core** | 92 | 13708 | 1,9 |
| **Total** | 4824 | 191283 | 100 |

**Table S3:** Core phylogeny and SNP matrix analysis of *G. vaginalis, G. swidsinskii, G. leopoldii* and *G. pickettii*.

| snp-dists 0.8.2 |  |  |  |  |  |  |  |  |  |  |  |  |  |
| --- | --- | --- | --- | --- | --- | --- | --- | --- | --- | --- | --- | --- | --- |
| *G. vaginalis* | c11Ua_71 | c15Ub_15 | c15Ub_3 | c15Ub_5 | c15Vsa_39 | c15Vsb_14 | c15Vsb_4 | c21Ua_54 | c29Ua_32 | c29Ub_7 | c29Vsa_19 | c29Vsb_3 |  |
| c11Ua_71 | 0 | 9922 | 9823 | 9977 | 9733 | 9806 | 9835 | 10119 | 9800 | 9804 | 9800 | 9804 | Vaginal swab |
| c15Ub_15 | 9922 | 0 | 1580 | 1275 | 518 | 1195 | 1572 | 7322 | 6972 | 6976 | 6972 | 6976 | Urine |
| c15Ub_3 | 9823 | 1580 | 0 | 665 | 1378 | 529 | 10 | 7374 | 7134 | 7138 | 7134 | 7138 |  |
| c15Ub_5 | 9977 | 1275 | 665 | 0 | 1379 | 786 | 673 | 7362 | 7181 | 7185 | 7181 | 7185 |  |
| c15Vsa_39 | 9733 | 518 | 1378 | 1379 | 0 | 1053 | 1370 | 7255 | 6962 | 6966 | 6962 | 6966 |  |
| c15Vsb_14 | 9806 | 1195 | 529 | 786 | 1053 | 0 | 523 | 7355 | 7077 | 7081 | 7077 | 7081 |  |
| c15Vsb_4 | 9835 | 1572 | 10 | 673 | 1370 | 523 | 0 | 7379 | 7139 | 7143 | 7139 | 7143 |  |
| c21Ua_54 | 10119 | 7322 | 7374 | 7362 | 7255 | 7355 | 7379 | 0 | 6436 | 6440 | 6436 | 6440 |  |
| c29Ua_32 | 9800 | 6972 | 7134 | 7181 | 6962 | 7077 | 7139 | 6436 | 0 | 4 | 0 | 4 |  |
| c29Ub_7 | 9804 | 6976 | 7138 | 7185 | 6966 | 7081 | 7143 | 6440 | 4 | 0 | 4 | 0 |  |
| c29Vsa_19 | 9800 | 6972 | 7134 | 7181 | 6962 | 7077 | 7139 | 6436 | 0 | 4 | 0 | 4 |  |
| c29Vsb_3 | 9804 | 6976 | 7138 | 7185 | 6966 | 7081 | 7143 | 6440 | 4 | 0 | 4 | 0 |  |
| *G. swidsinskii* |  | c15Ua_15 | c15Ua_23 | c15Ub_13 | c15Vsa_36 | c15Vsb_13 |  |  |  |  |  |  |  |
| c10Ua_244 | 0 | 9196 | 9201 | 9201 | 9197 | 9201 |  |  |  |  |  |  |  |
| c15Ua_15 | 9196 | 0 | 11 | 11 | 3 | 11 |  |  |  |  |  |  |  |
| c15Ua_23 | 9201 | 11 | 0 | 2 | 12 | 2 |  |  |  |  |  |  |  |
| c15Ub_13 | 9201 | 11 | 2 | 0 | 12 | 0 |  |  |  |  |  |  |  |
| c15Vsa_36 | 9197 | 3 | 12 | 12 | 0 | 12 |  |  |  |  |  |  |  |
| c15Vsb_13 | 9201 | 11 | 2 | 0 | 12 | 0 |  |  |  |  |  |  |  |
| *G. leopoldii* | c18Ua_11<br>0 | c18Ua_112 | c1Ub_40 | c1Ub_6 | c31Ua_40 | c31VSa_14 | c31VSa_35 |  |  |  |  |  |  |
| c18Ua_110 | 0 | 0 | 8006 | 8006 | 7737 | 7737 | 7735 |  |  |  |  |  |  |
| c18Ua_112 | 0 | 0 | 8006 | 8006 | 7737 | 7737 | 7735 |  |  |  |  |  |  |
| c1Ub_40 | 8006 | 8006 | 0 | 3 | 7257 | 7257 | 7261 |  |  |  |  |  |  |
| c1Ub_6 | 8006 | 8006 | 3 | 0 | 7256 | 7256 | 7259 |  |  |  |  |  |  |
| c31Ua_40 | 7737 | 7737 | 7257 | 7256 | 0 | 0 | 0 |  |  |  |  |  |  |
| c31VSa_14 | 7737 | 7737 | 7257 | 7256 | 0 | 0 | 0 |  |  |  |  |  |  |
| c31VSa_35 | 7735 | 7735 | 7261 | 7259 | 0 | 0 | 0 |  |  |  |  |  |  |
| *G. pickettii* | 10Ua_31 | 12Ub_1 | 17Ua_112 | 23Ua_15 | 23Ua_38 | 17Ua_118 | 23Ub_4 |  | Asymptomatic |  |  |  |  |
| c10Ua_31 | 0 | 19186 | 18348 | 18348 | 18348 | 19875 | 18538 |  | OAB |  |  |  |  |
| c12Ub_1 | 19186 | 0 | 18704 | 18704 | 18704 | 7707 | 18318 |  |  |  |  |  |  |
| c17Ua_112 | 18348 | 18704 | 0 | 1 | 1 | 20162 | 18683 |  |  |  |  |  |  |
| c23Ua_15 | 18348 | 18704 | 1 | 0 | 1 | 20162 | 18683 |  |  |  |  |  |  |
| c23Ua_38 | 18348 | 18704 | 1 | 1 | 0 | 20162 | 18683 |  |  |  |  |  |  |
| c17Ua_118 | 19875 | 7707 | 20162 | 20162 | 20162 | 0 | 19902 |  |  |  |  |  |  |
| c23Ub_4 | 18538 | 18318 | 18683 | 18683 | 18683 | 19902 | 0 |  |  |  |  |  |  |

**Table S4:**
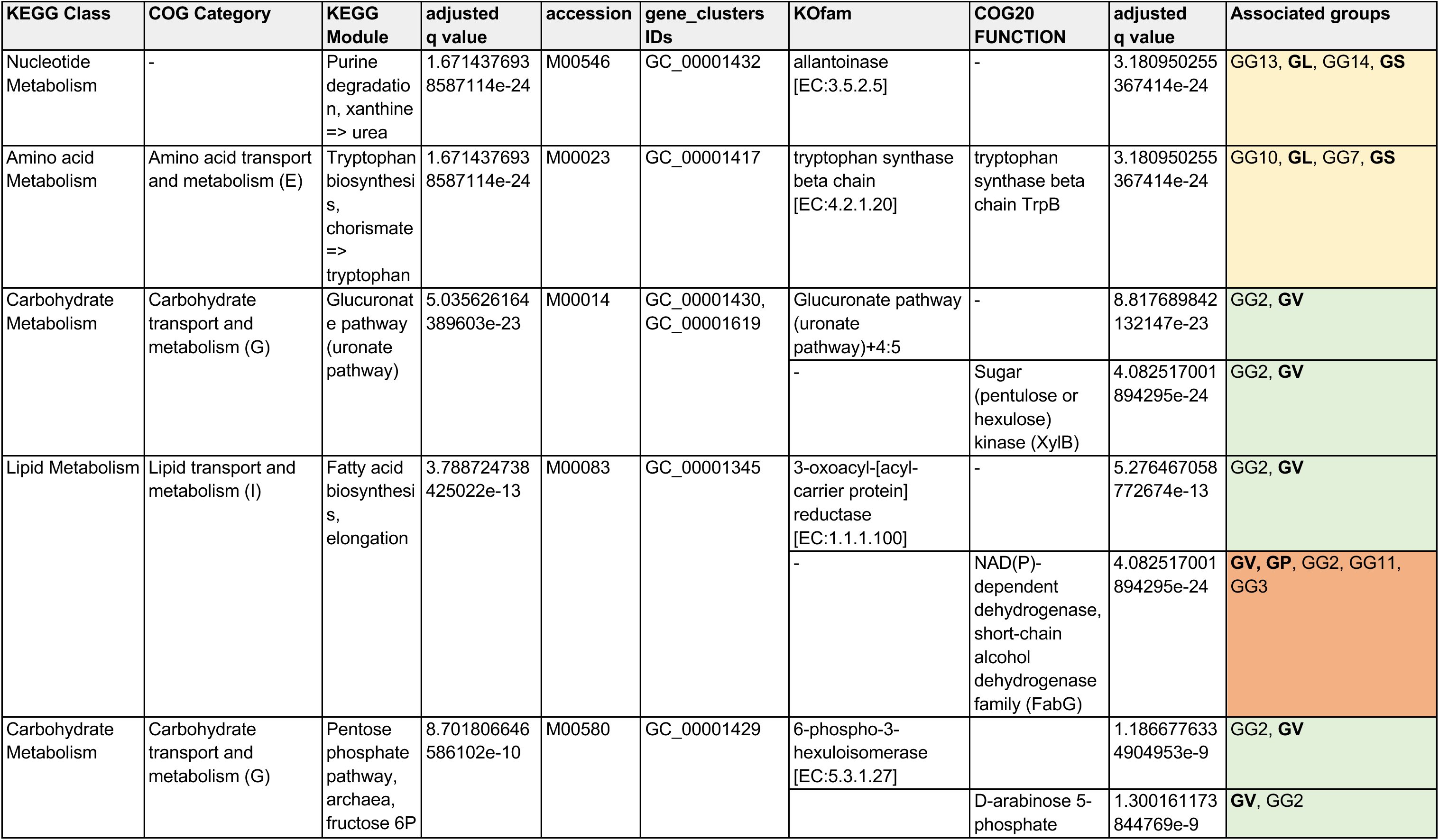
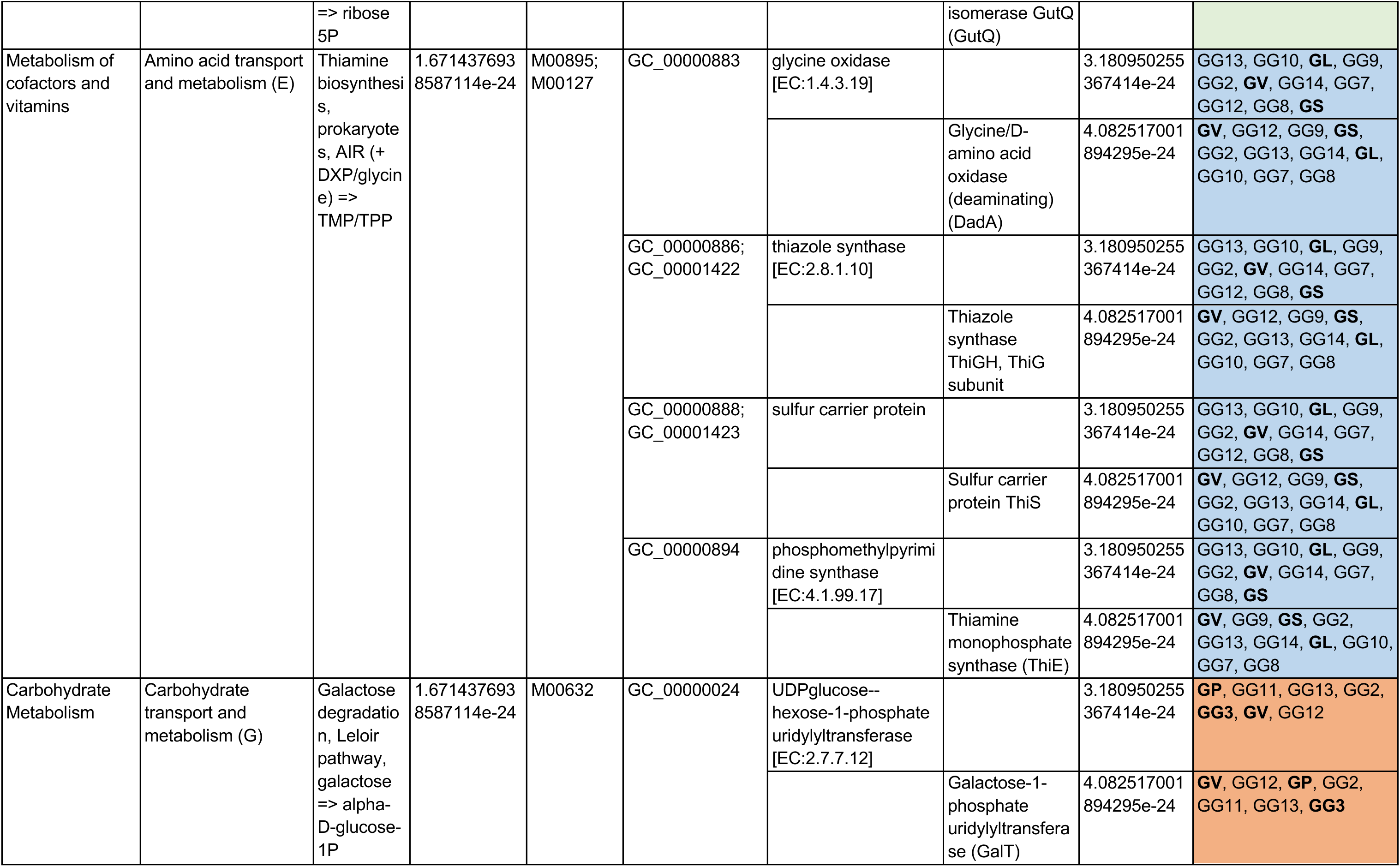
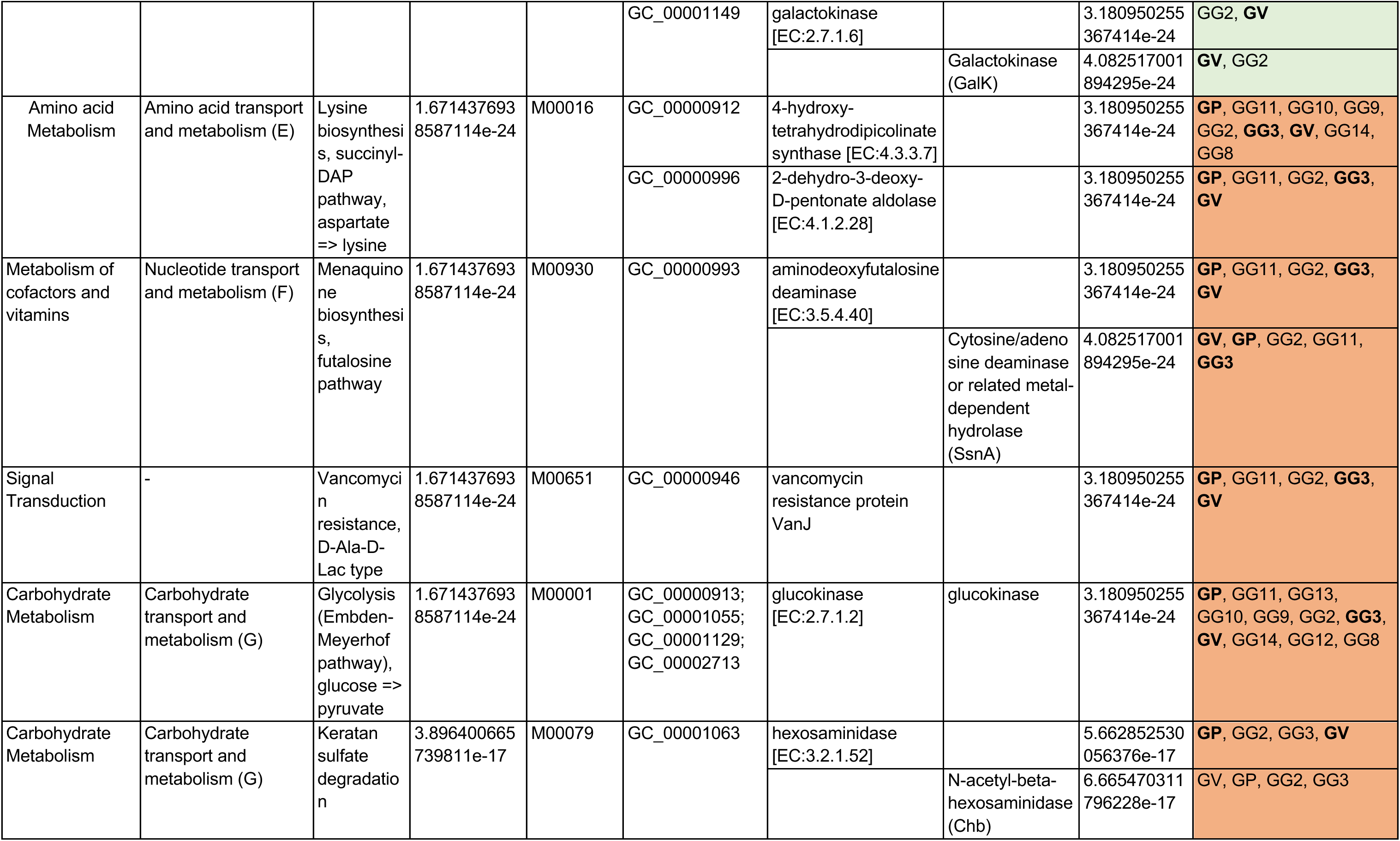
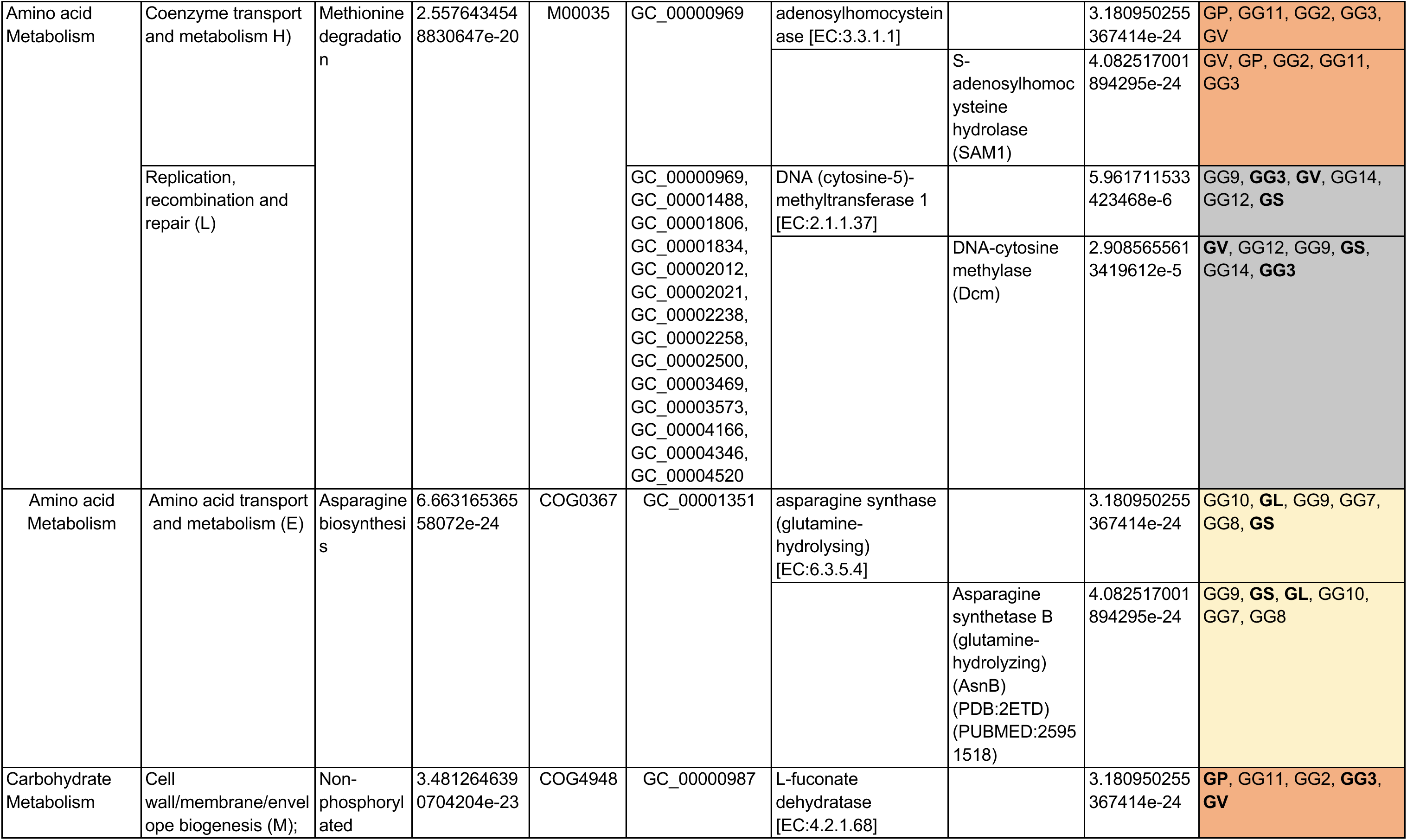
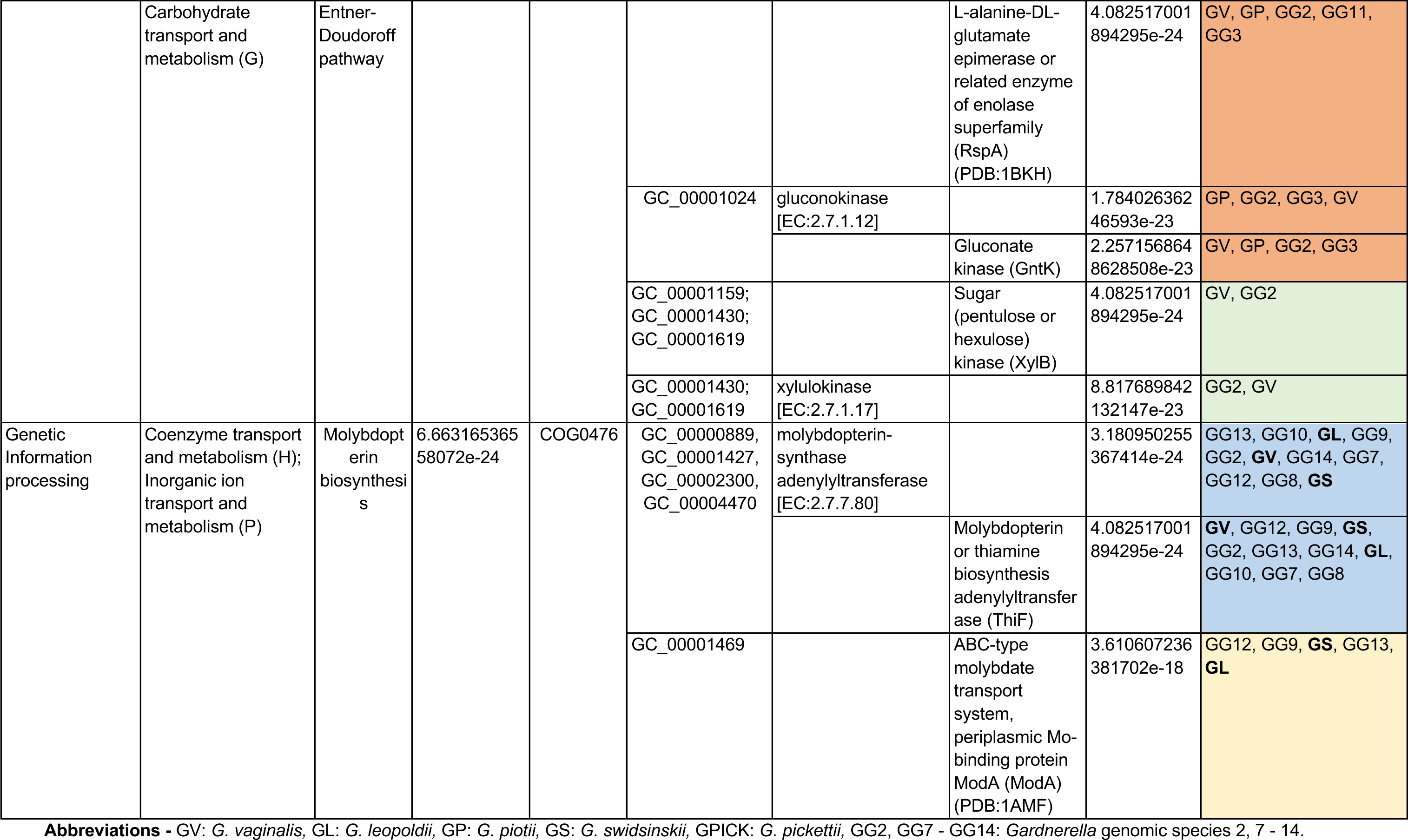
Functional enrichment analysis through KEGG modules and COG pathways enriched in specific *Gardnerella* species.

**Table S5:**
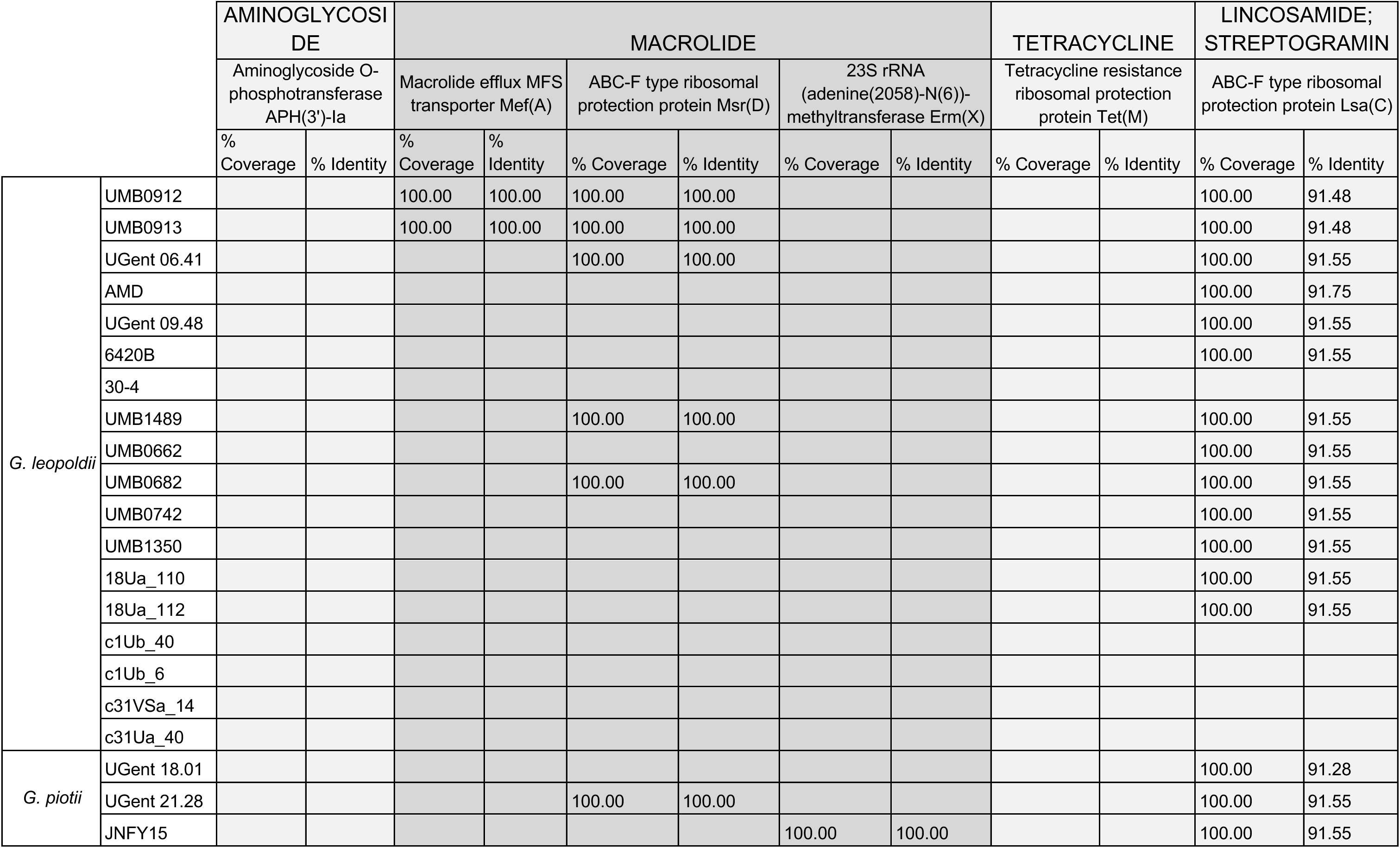
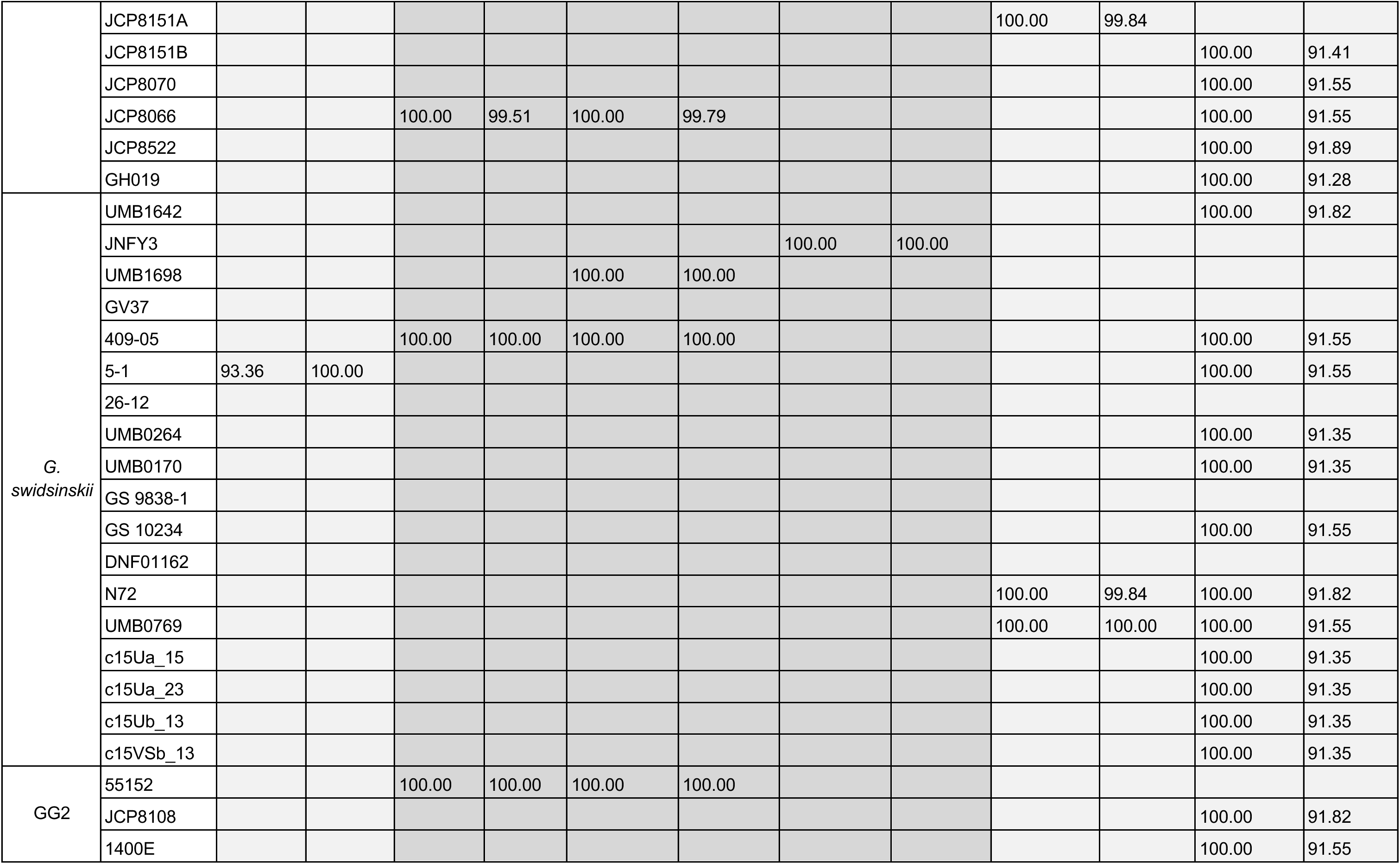
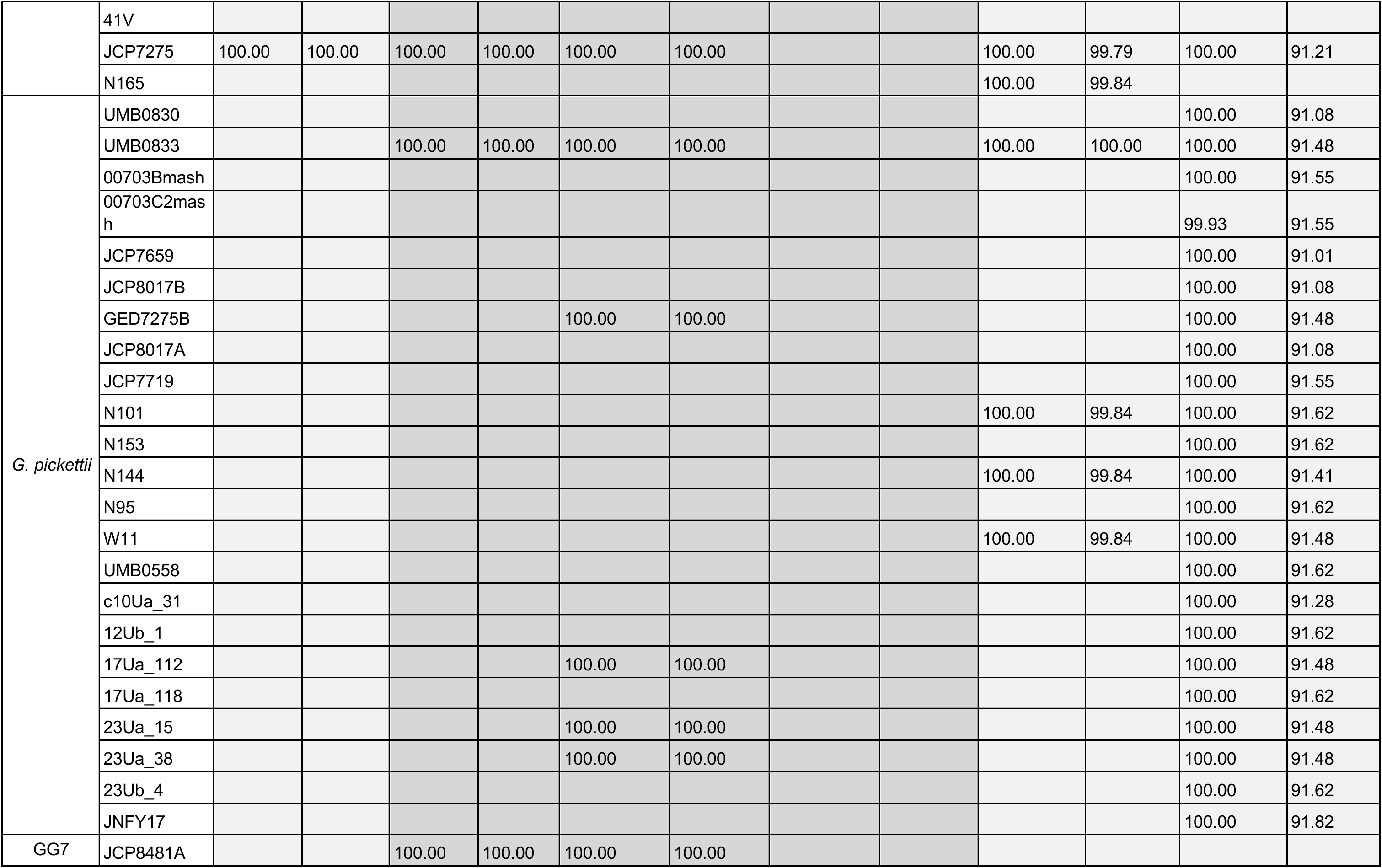
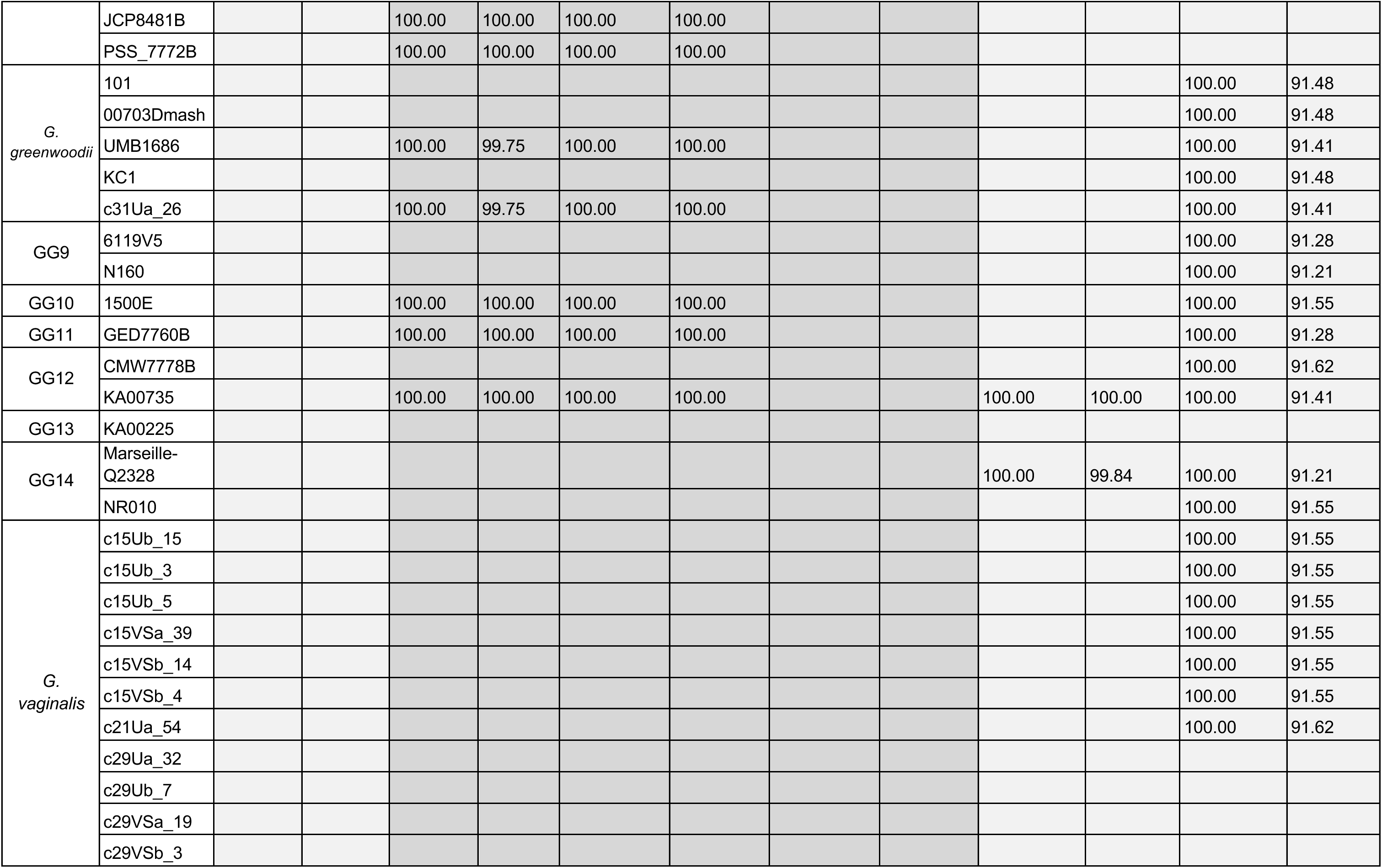
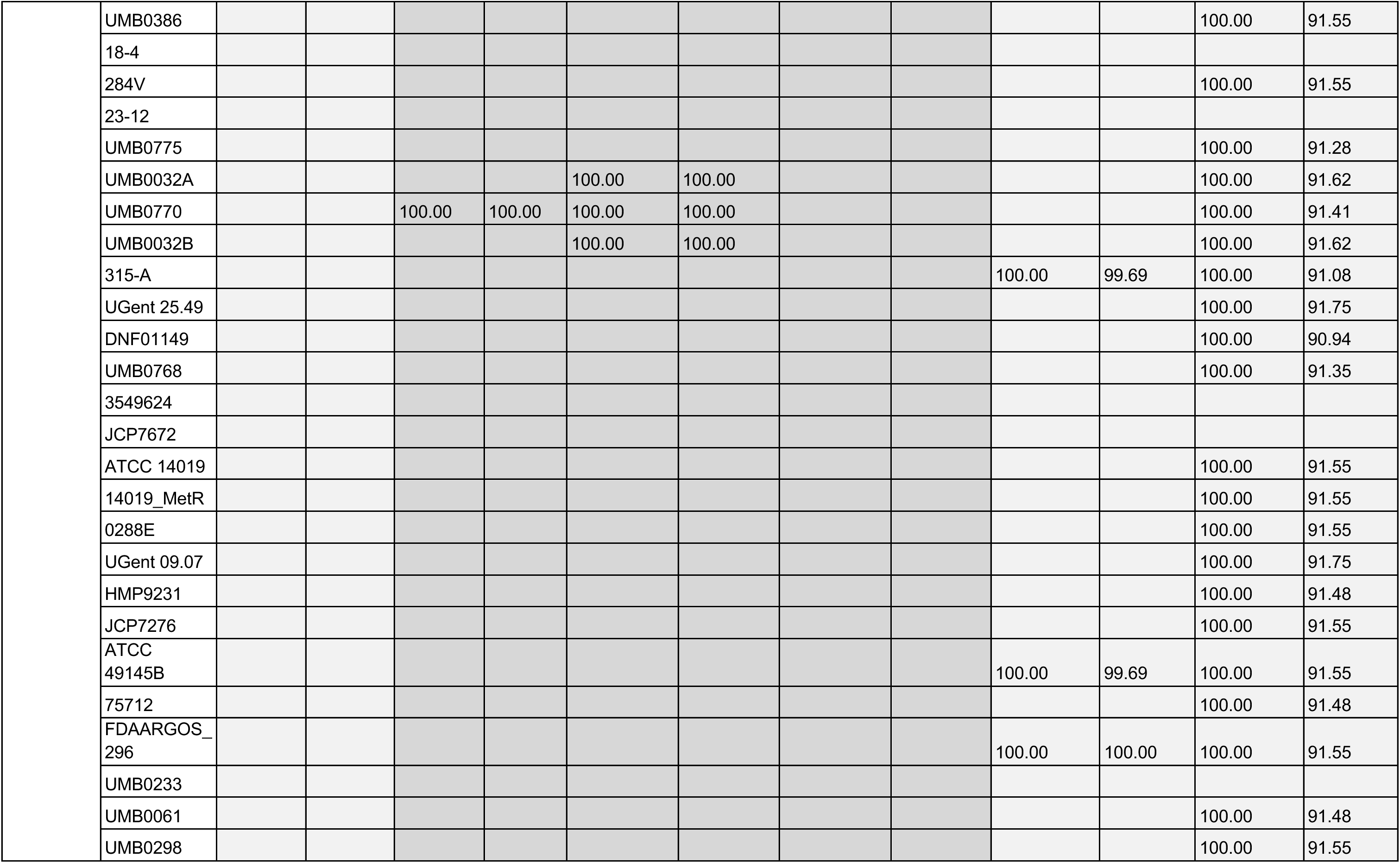
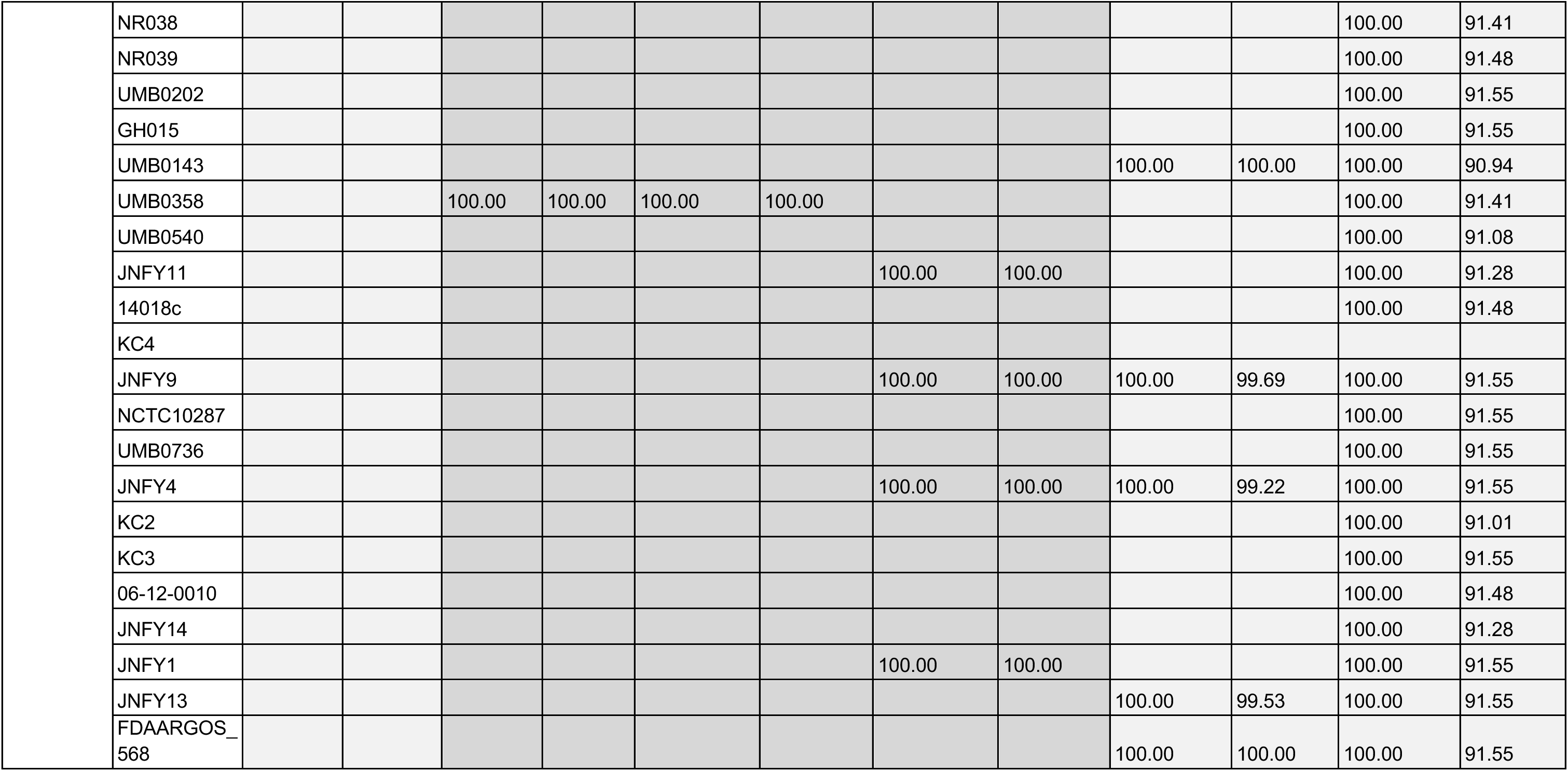
Antibiotic Resistance Genes identified in *Gardnerella* pangenome.

**Table S6:**
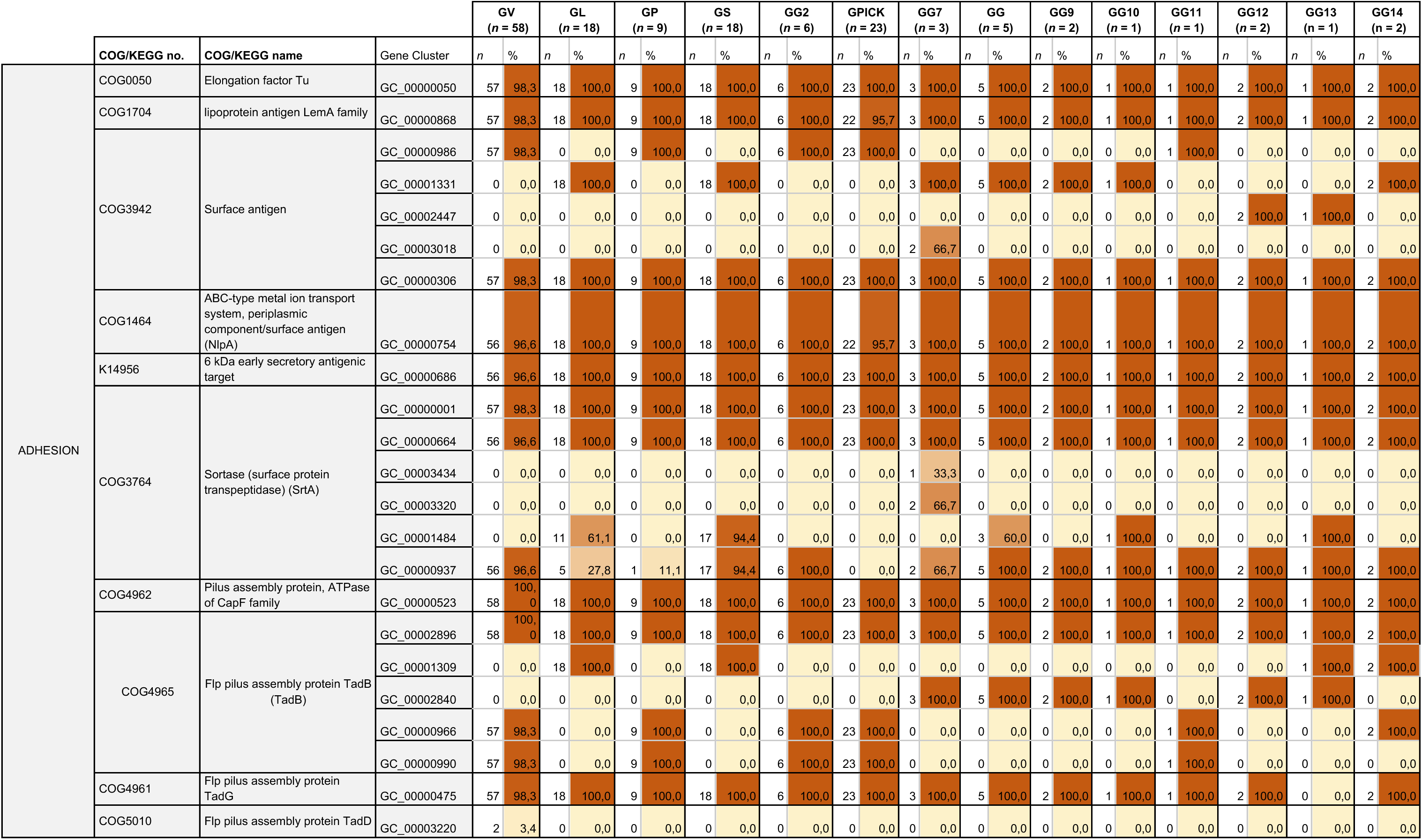
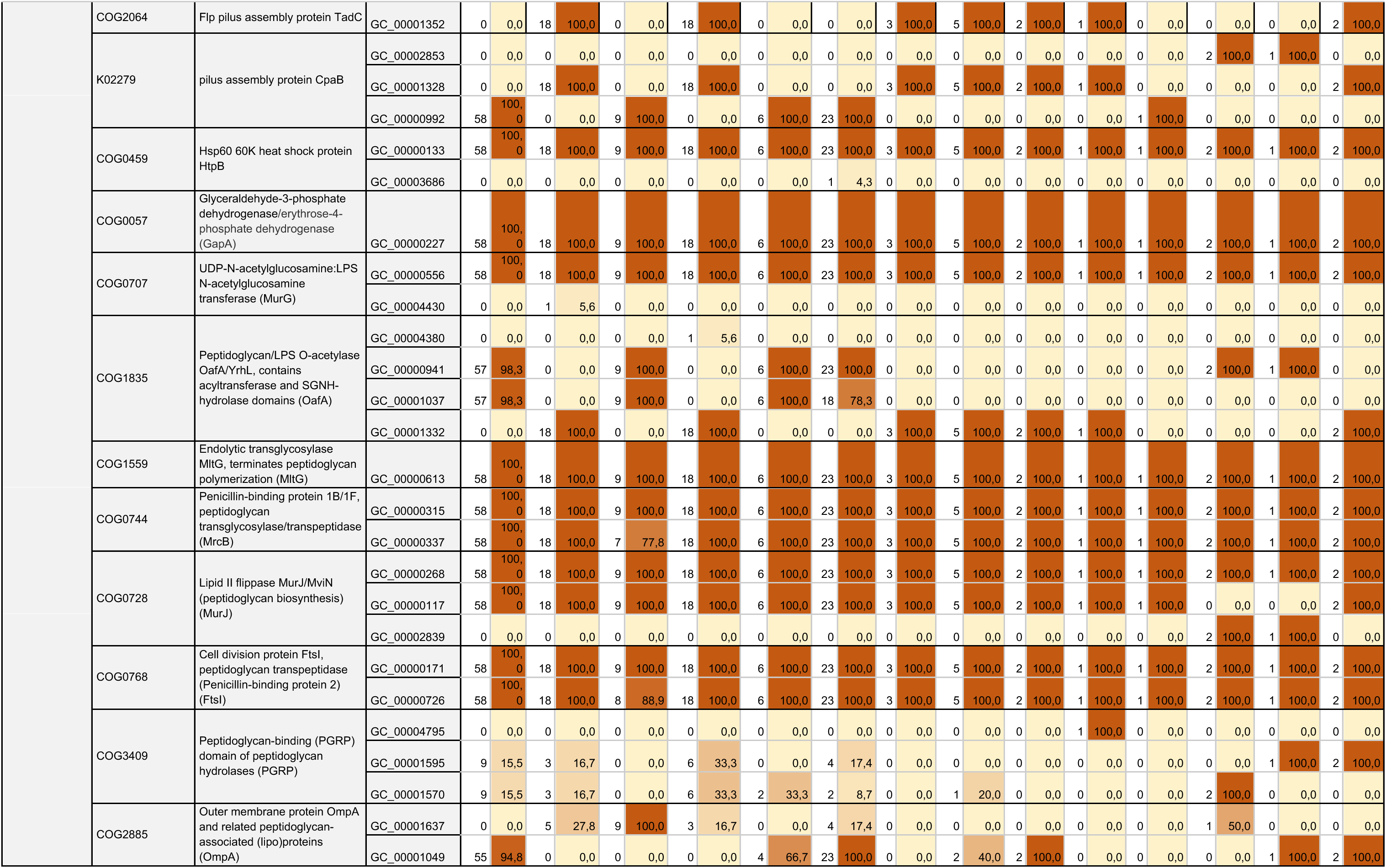
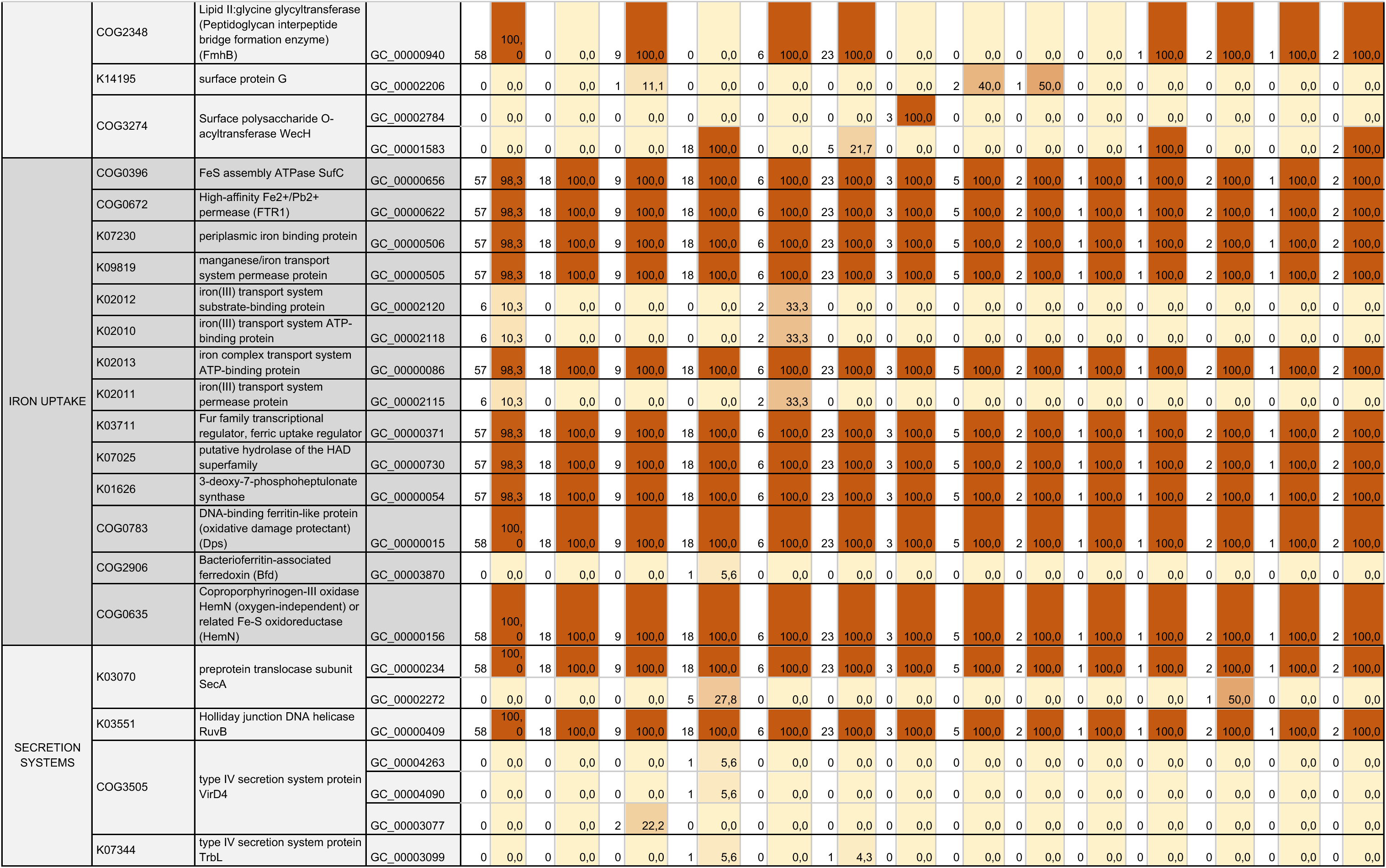
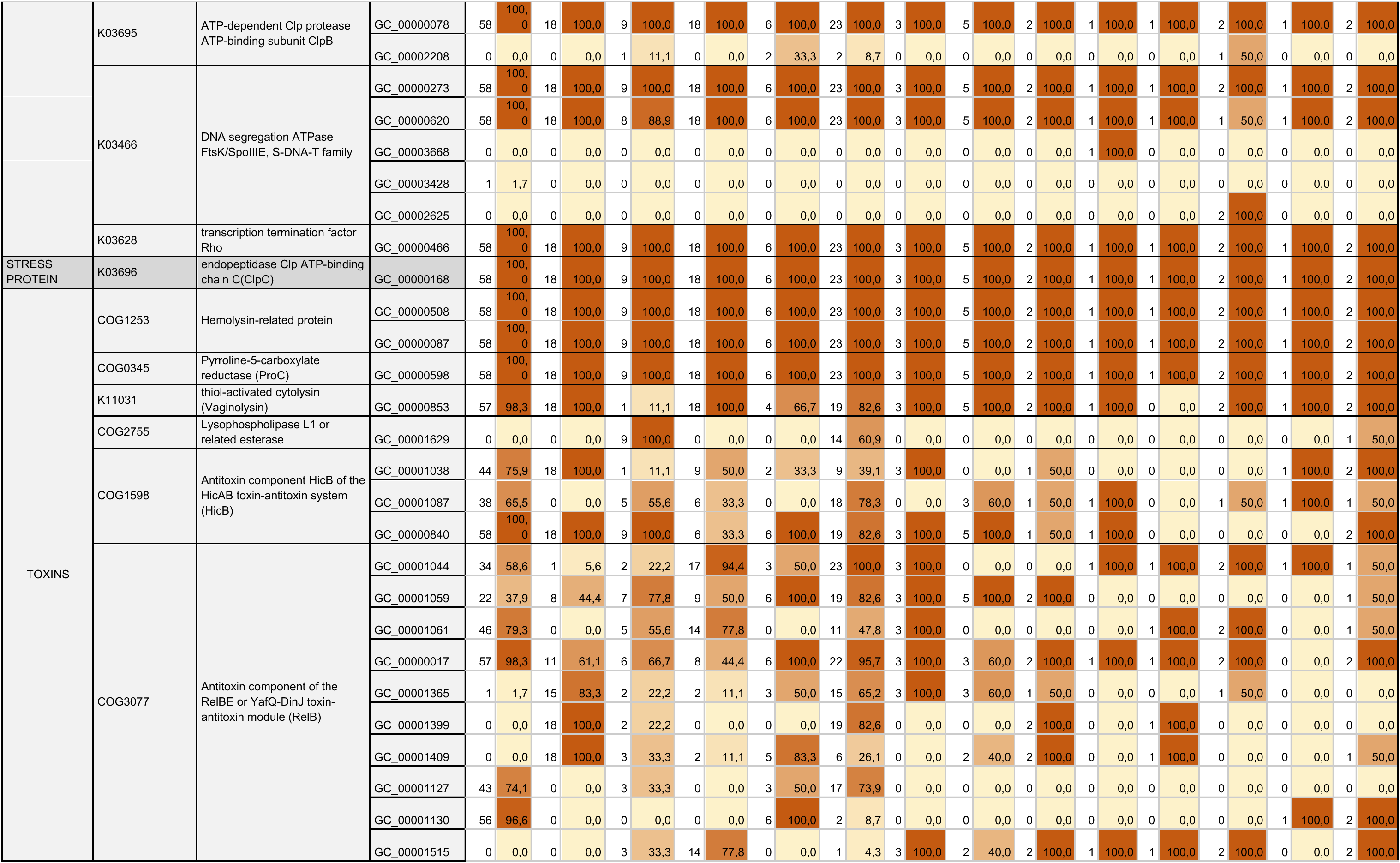
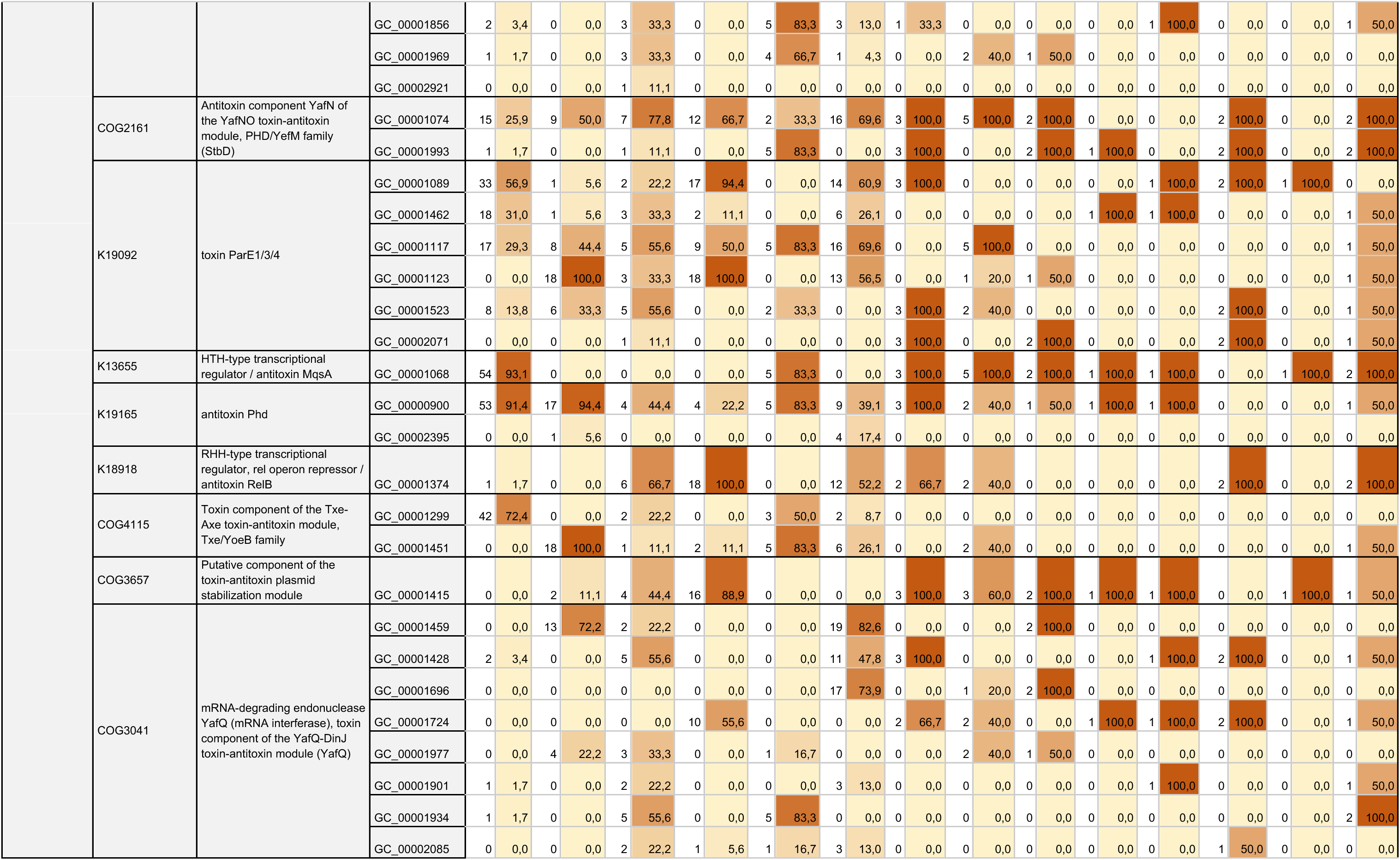
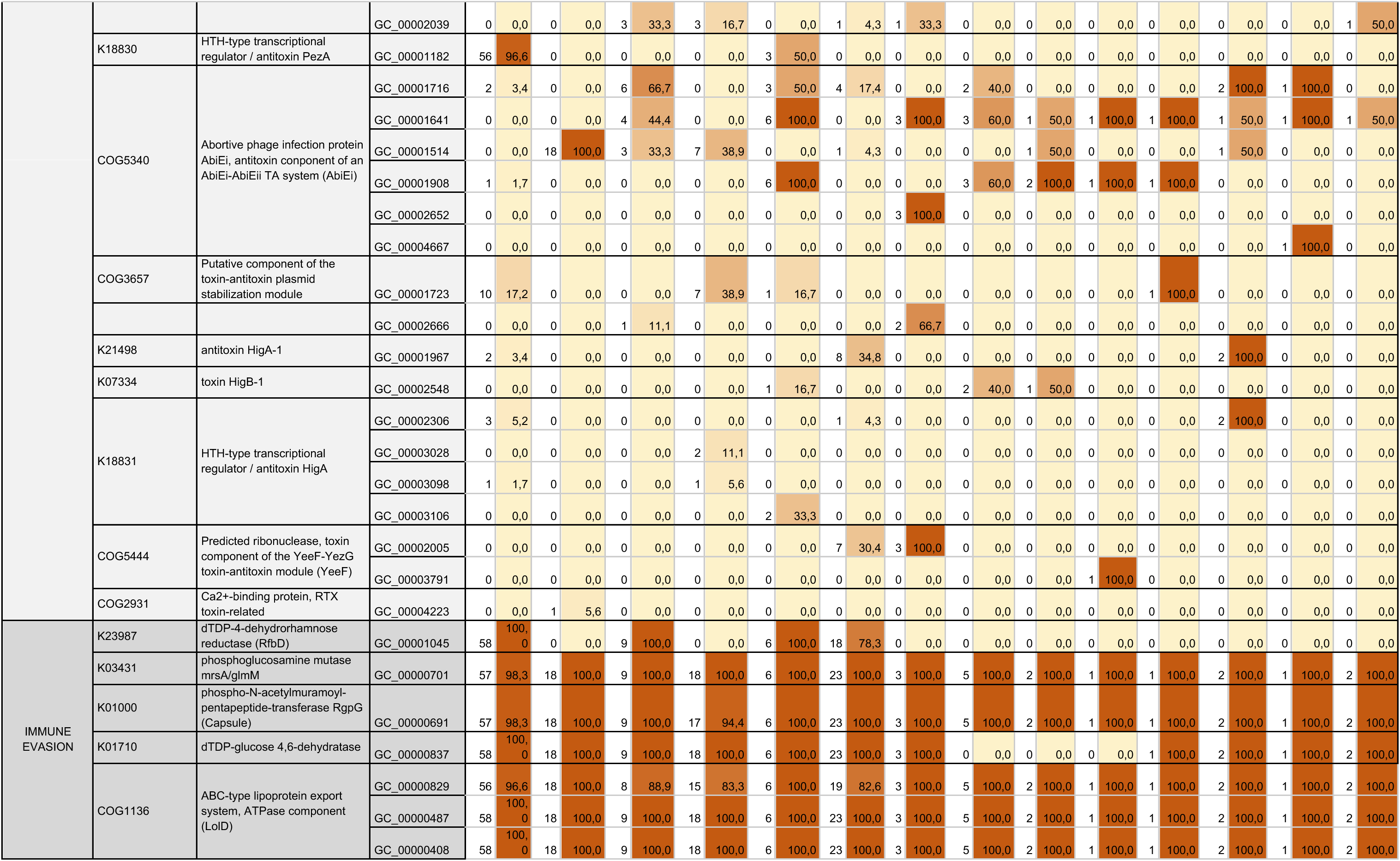
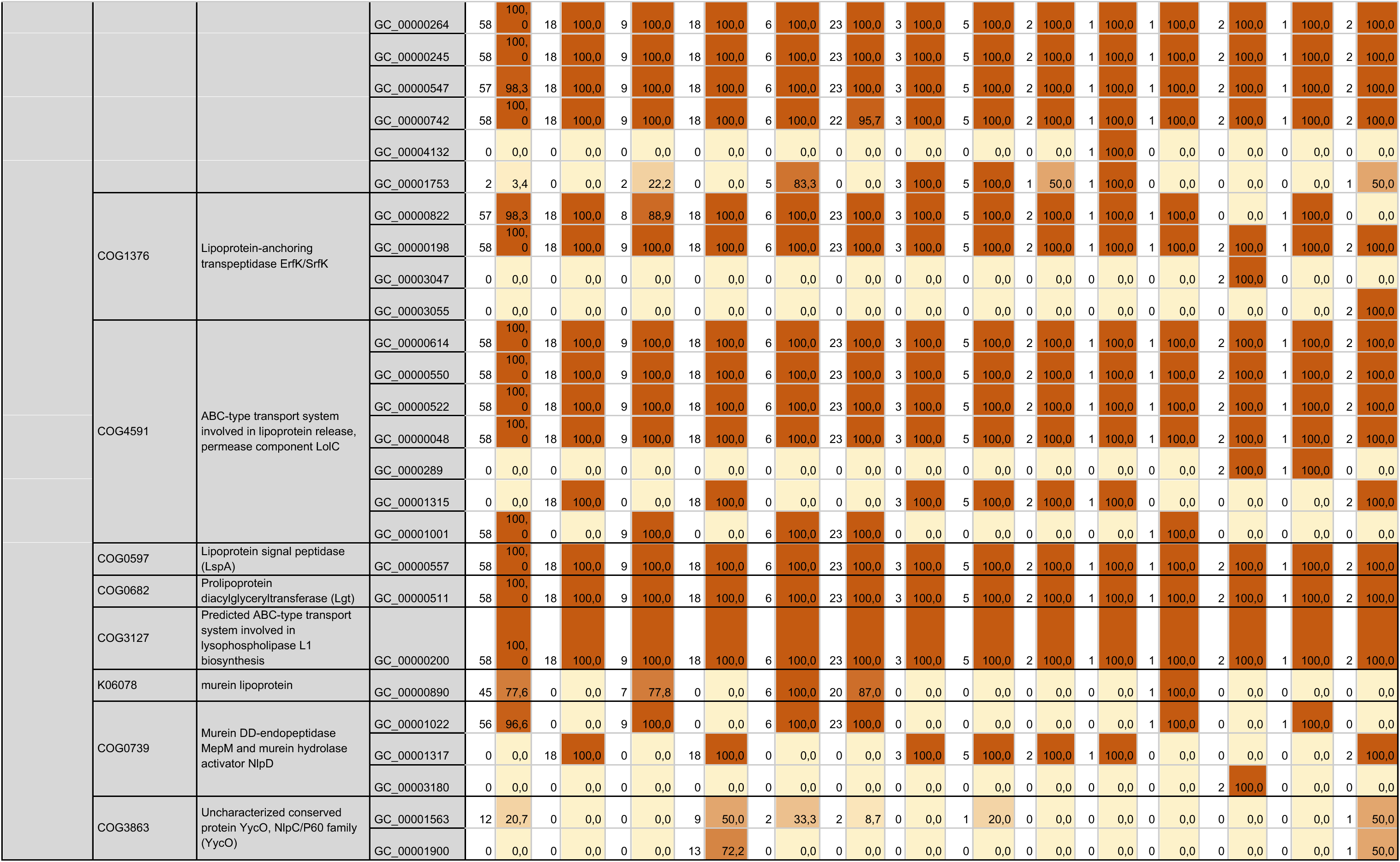
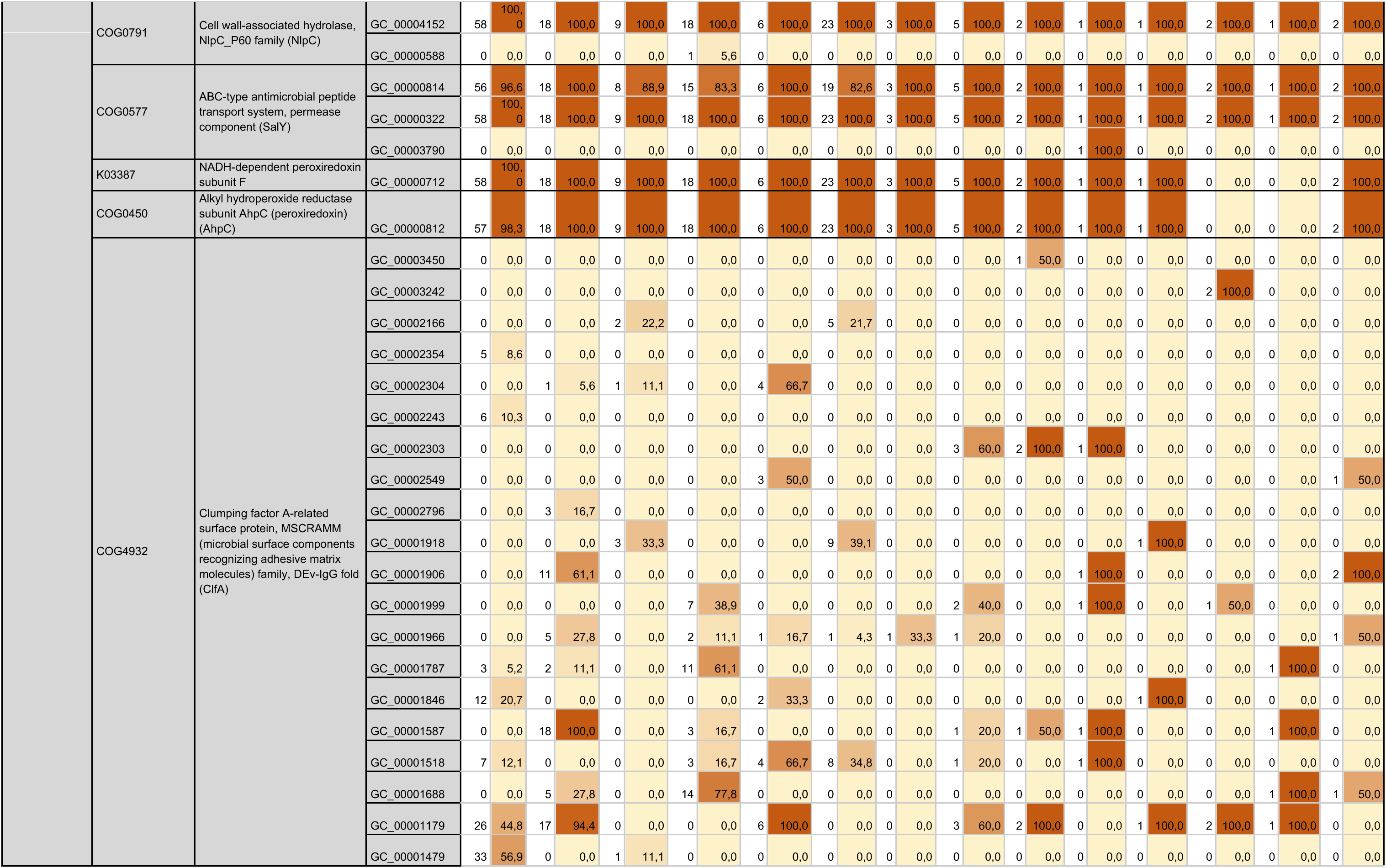
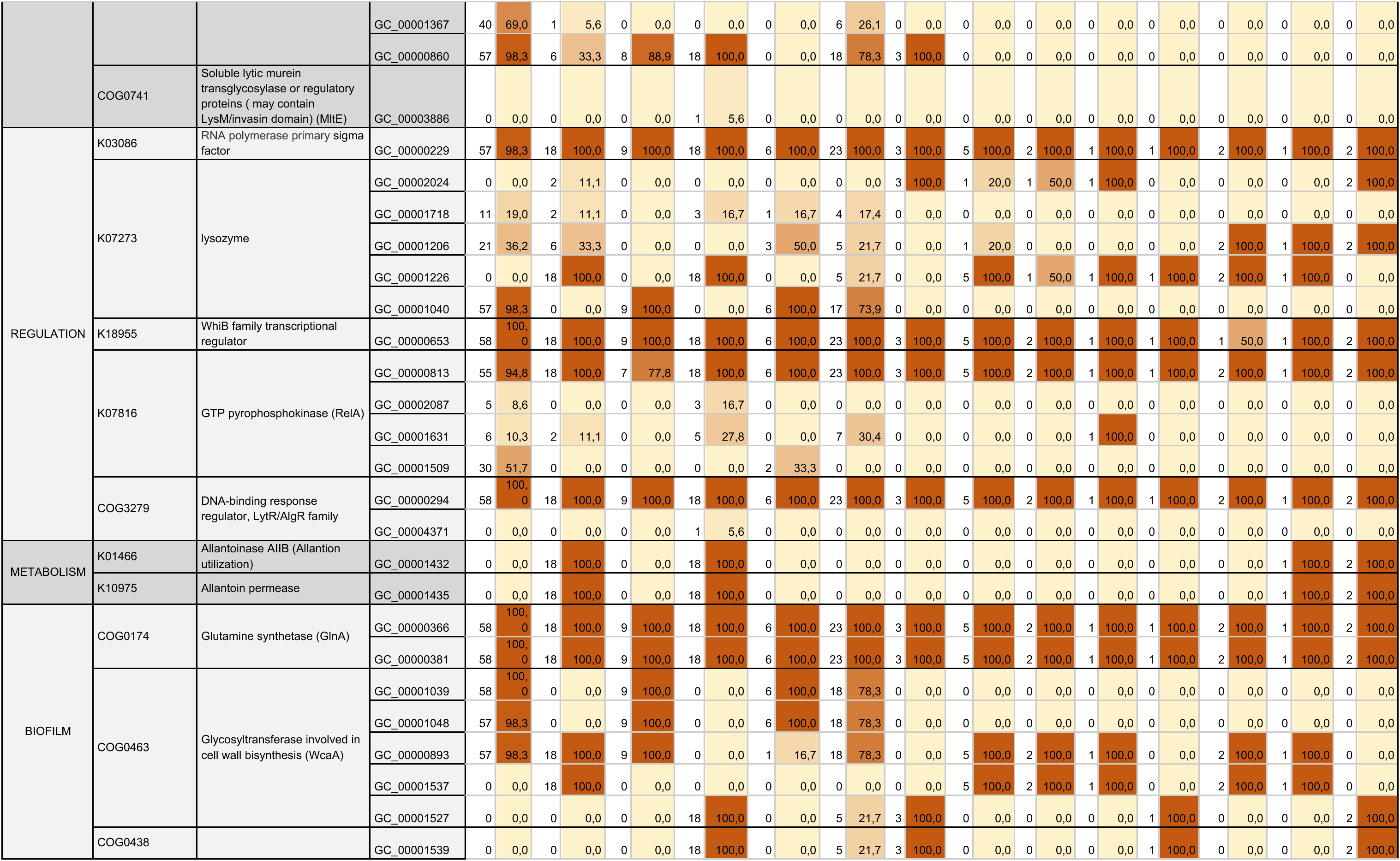
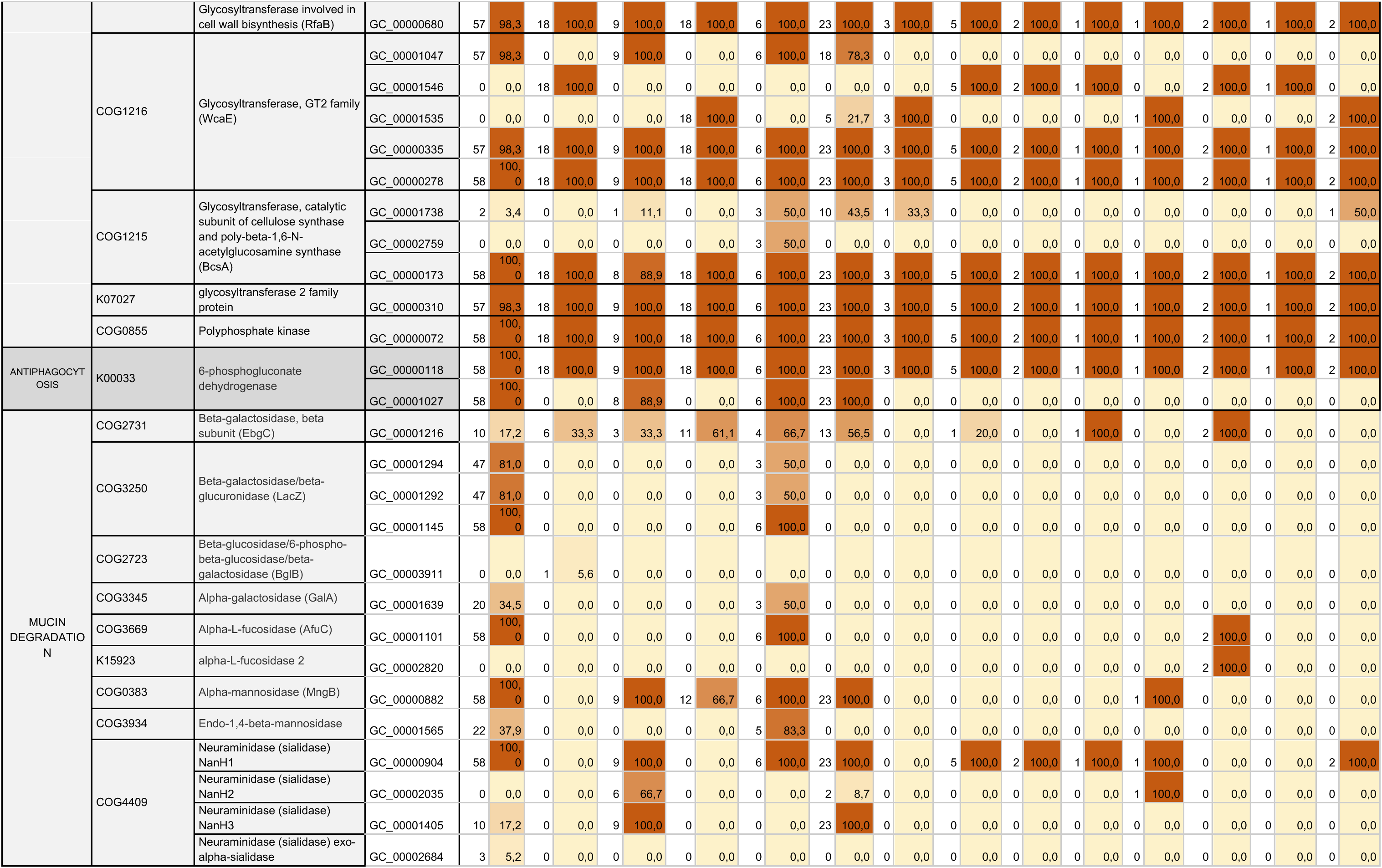
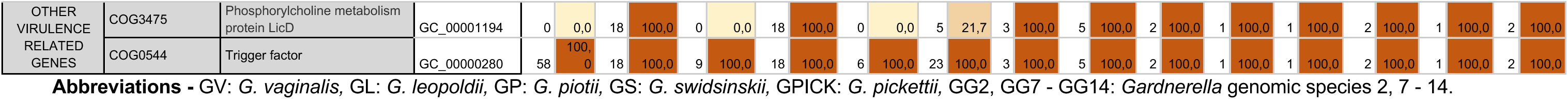
Virulence-associated functions and gene clusters in the *Gardnerella* genus.

**Table S7:** Secondary metabolite biosynthetic gene clusters (BGCs) across *Gardnerella* genomes.

|  | GV<br>(n = 58) |  | GL<br>(n = 18) |  | GP<br>(n = 9) |  | GS<br>(n = 18) |  | GG2<br>(n = 6) |  | GPICK<br>(n = 23) |  | GG7<br>(n = 3) |  | GG<br>(n = 5) |  | GG9<br>(n = 2) |  | GG10<br>(n = 1) |  | GG11<br>(n = 1) |  | GG12<br>(n = 2) |  | GG13<br>(n = 1) |  | GG14<br>(n = 2) |  |
| --- | --- | --- | --- | --- | --- | --- | --- | --- | --- | --- | --- | --- | --- | --- | --- | --- | --- | --- | --- | --- | --- | --- | --- | --- | --- | --- | --- | --- |
|  | n | % | n | % | n | % | n | % | n | % | n | % | n | % | n | % | n | % | n | % | n | % | n | % | n | % | n | % |
| Class II lanthipeptide clusters like mutacin II (U40620) | 0 | 0,0 | 0 | 0,0 | 0 | 0,0 | 15 | 83,3 | 0 | 0,0 | 0 | 0,0 | 0 | 0,0 | 0 | 0,0 | 0 | 0,0 | 0 | 0,0 | 0 | 0,0 | 0 | 0,0 | 0 | 0,0 | 0 | 0,0 |
| Other unspecified ribosomally synthesised and post-translationally modified peptide product (RiPP) cluster | 1 | 1,7 | 0 | 0,0 | 0 | 0,0 | 18 | 100,0 | 0 | 0,0 | 0 | 0,0 | 0 | 0,0 | 0 | 0,0 | 0 | 0,0 | 0 | 0,0 | 0 | 0,0 | 0 | 0,0 | 0 | 0,0 | 0 | 0,0 |
| Type III PKS | 54 | 93,1 | 1 | 5,6 | 4 | 44,4 | 0 | 0,0 | 3 | 50,0 | 19 | 82,6 | 0 | 0,0 | 0 | 0,0 | 0 | 0,0 | 0 | 0,0 | 0 | 0,0 | 2 | 100,0 | 0 | 0,0 | 0 | 0,0 |
| Linear azol(in)e-containing peptides | 2 | 3,4 | 0 | 0,0 | 0 | 0,0 | 1 | 5,6 | 0 | 0,0 | 0 | 0,0 | 0 | 0,0 | 0 | 0,0 | 0 | 0,0 | 0 | 0,0 | 0 | 0,0 | 0 | 0,0 | 0 | 0,0 | 0 | 0,0 |

**Abbreviations** - GV: *G. vaginalis*, GL: *G. leopoldii*, GP: *G. piotii*, GS: *G. swidsinskii*, GPICK: *G. pickettii*, GG2, GG7 - GG14: *Gardnerella* genomic species 2, 7 - 14.

